# A nanobody-based proximity ligation assay detects constitutive and stimulus-regulated native Arc/Arg3.1 oligomers in hippocampal neuronal dendrites

**DOI:** 10.1101/2024.08.14.607905

**Authors:** Rodolfo Baldinotti, Francois P. Pauzin, Hauk Fevang, Yuta Ishizuka, Clive R. Bramham

**Affiliations:** Department of Biomedicine, University of Bergen, Bergen, Norway; Mohn Research Center for the Brain, University of Bergen, Norway

**Keywords:** Arc protein, nanobody, proximity ligation assay, oligomerization, synaptic plasticity hippocampus

## Abstract

Activity-regulated cytoskeleton-associated (Arc), the product of an immediate early gene, plays critical roles in synaptic plasticity and memory. Evidence suggests that Arc function is determined by its oligomeric state, however, methods for localization of native Arc oligomers are lacking. Here, we developed a nanobody-based proximity ligation assay (PLA) for detection, localization, and quantification of Arc-Arc complexes in primary rat hippocampal neuronal cultures. We used nanobodies with single, structurally defined epitopes in the bilobar Arc capsid domain. Nanobody H11 binds inside the N-lobe ligand pocket, while nanobody C11 binds to the C-lobe surface. For each nanobody, ALFA- and FLAG-epitope tags created a platform for antibody binding and PLA. Surprisingly, PLA puncta in neuronal dendrites revealed widespread constitutive Arc-Arc complexes. Treatment of cultures with tetrodotoxin or cycloheximide had no effect, suggesting stable complexes that are independent of recent neuronal activity and protein synthesis. To assess detection of oligomers, cultures were exposed to a cell-penetrating peptide inhibitor of the Arc oligomerization motif (OligoOFF). Arc-Arc complexes detected by H11 PLA were inhibited by OligoOff but not by control peptide. Notably, Arc complexes detected by C11 were unaffected by OligoOFF. Furthermore, we evaluated Arc complex formation after chemical stimuli that increase Arc synthesis. Brain-derived neurotrophic factor increased Arc-Arc signal detected by C11, but not H11. Conversely, dihydroxyphenylglycine (DHPG) treatment selectively enhanced H11 PLA signals. In sum, nanobody-based PLA reveals constitutive and stimulus-regulated Arc oligomers in hippocampal neuronal dendrites. A model is proposed based on detection of Arc dimer by C11 and higher-order oligomer by H11 nanobody.

## Introduction

Understanding how the brain encodes information for learning and memory is one of the greatest challenges in neuroscience. Representation of information in neural networks depends on synaptic plasticity mechanisms; the ability of synapses to undergo changes in the efficacy of synaptic transmission [1–3]. For long-term modifications (hours to days), *de novo* gene transcription is required [4]. Activity-regulated cytoskeleton-associated protein (Arc; or activity-regulated gene, Arg3.1), regulates several forms of plasticity at excitatory synapses, such as long-term potentiation (LTP), long-term depression (LTD), and synaptic scaling, and functions in postnatal cortical development and memory formation [5]. Following stimulus-evoked transcription, Arc mRNA is found in cell body and dendrites and accumulates at stimulated post-synaptic sites for local translation. Depending on the synaptic stimulation pattern, newly synthesized Arc acts transiently to support the formation of LTP or LTD [6–10], with an estimated protein half-life of 30 to 60 minutes [11–13]. Exactly how Arc is targeted toward specific types of plasticity and cellular functions is not fully understood. However, mounting evidence implicates Arc oligomeric state as a major determinant of its function [14–17].

Arc evolved from ancient Ty3/Gypsy retrotransposon elements and has conserved homology with Group-specific antigen (Gag) polyprotein in modern retroviruses [18, 19]. A salient feature of purified mammalian Arc protein is its ability to self-assemble into stable low and high-order oligomeric species (dimer to 32mer) *in vitro* [20–23]. Mice harboring point mutations in residues important for Arc oligomerization exhibit altered synaptic plasticity and memory [22]. The largest oligomeric form of Arc is a retrovirus-like capsid containing Arc mRNA [24, 25]. In neuronal cultures, Arc capsids are released from neurons in extracellular vesicles, which transmit the capsid and release the RNA for expression of Arc in neighboring cells [24, 26, 27]. Using *in situ* gene labeling, evidence was obtained for transfer of tagged Arc protein between neurons in adult mouse hippocampus [28]. Recent advances in imaging Arc oligomers are based on ectopic expression of tagged protein in non-neuronal cells [29–31]. However, little is known about the formation and regulation of native Arc oligomers. In situ protein crosslinking of rat brain regions identified an Arc dimer with low constitutive expression and enhanced expression after in vivo LTP induction or treatment of hippocampal neuronal cultures with brain-derived neurotrophic factor (BDNF) [32]. Thus, to elucidate Arc function, new strategies are needed for detection, localization, and quantification of native oligomers.

Mammalian Arc has as N-terminal coiled-coil domain and a C-terminal capsid (CA) domain separated by a flexible linker [14]. The CA has two lobes, the N-lobe and C-lobe, with 3D structural homology to the retroviral Gag CA domain which mediates virion capsid assembly [19, 33]. Uniquely, the mammalian Arc N-lobe harbours a binding pocket for several synaptic protein ligands [19]. Single-domain antibodies, also known as nanobodies, are 10 times smaller than conventional antibodies with unique binding properties [34, 35]. Recently, we developed six anti-Arc nanobodies (clones B5, B12, C11, D4, E5 and H11) from immunized Alpaca and further validated application of the recombinant ALFA epitope-tagged nanobodies for immunocytochemistry and immunoprecipitation [36, 37]. Epitope mapping and isothermal calorimetry assays showed that all six nanobodies bind to Arc CA with nanomolar affinity [36, 37]. Furthermore, crystal structure analysis showed that nanobody H11 binds specifically to the N-lobe ligand pocket [37, 38].

Here, we exploited the specific epitope binding anti-Arc nanobodies to develop a novel proximity ligation assay (PLA) for detection of native Arc-Arc complexes. PLA detects protein-protein proximity *in situ*, amplifying the signal to give cellular localization of complexes that might be present in low levels [39–41]. Using H11 fused to ALFA and FLAG epitope tags, we identify constitutive Arc-Arc complexes in neuronal dendrites of unstimulated hippocampal neuronal cultures. We provide evidence that these complexes contain stable oligomers that do not require recent neuronal activity or *de novo* Arc expression. Constitutive Arc-Arc complexes are similarly detected by PLA using nanobody C11, which binds to an exposed surface of the capsid domain C-lobe [37]. Comparing chemical stimuli that enhance Arc protein expression, we demonstrate stimulus-evoked changes in Arc-Arc PLA consistent with formation of dimers and higher-order oligomers.

## Material and methods

### Primary hippocampal neuronal culture preparation

All animal procedures used for the primary culture preparation were performed accordingly to the Norwegian animal care committee established regulations. This research is approved by the Norwegian National Research Ethics Committee in compliance with EU Directive 2010/63/EU, ARRIVE guidelines. The neuronal primary culture preparation was based on the protocols published for neuronal culture preparation and cryopreservation by Ishizuka et al., 2020 [42]. Before the embryonic dissection for the hippocampus isolation, coverslips (Marienfeld-Superior, Lauda-Königshofen, Germany) of 12 or 18 mm were treated with a 1 N HNO_3_ solution (Sigma-Aldrich, St. Louis, MO), followed by washing step with Mili-Q® sterile water and coverslip coating with 1 mg/mL poly-l-lysine (PLL; Sigma-Aldrich) in borate buffer (Thermo Fisher Scientific, Waltham, MA).

Briefly, for the animal dissection and cell culture preparation, pregnant Wistar rats were deeply anesthetized and submitted to euthanasia by CO_2_. Embryos from embryonic days 18 (E18) were removed from the uterus and micro dissected for hippocampal collection, performed in cold Hank’s Balanced Salt Solution (HBSS), containing 10 mM Hepes (Thermo Fisher Scientific) and 100 U/mL Penicillin-Streptomycin under a laminar air-flow hood. After isolation, the hippocampi were kept in cold HEPES buffer, followed by tissue and cellular dissociation (37 °C), using 0.05% Trypsin-EDTA solution (Thermo Fisher Scientific) containing 10 mM HEPES and 100 U/mL Penicillin-Streptomycin (Thermo Fisher Scientific. After trypsin treatment, the tissue was mechanically dissociated by repetitive pipetting with a flame-polished glass Pasteur pipette. The cells were counted in a Neubauer chamber and were either cryopreserved in freezing media (80 % FBS + 20 % DMSO) or seeded in culture plates.

Fresh or cryopreserved neurons were seeded in a cell density of 30.000 or 60.000 cells/well (24-well plate and 12-well plate, respectively) in cell-culture plates containing pre-coated coverslips in Minimum Essential Medium (MEM, Thermo Fisher Scientific), supplemented with 10% fetal bovine serum (FBS) (Sigma-Aldrich), 0.6% glucose (Sigma-Aldrich), and 1 mM sodium pyruvate (Thermo Fisher Scientific), and 100 U/ml Penicillin-Streptomycin). After the cells attached, the platting medium was replaced by Neurobasal™ Medium (Thermo Fisher Scientific), supplemented with 2% B-27™ (Thermo Fisher Scientific) and 0.25% GlutaMAX™-I (Thermo Fisher Scientific). On days *in vitro* (DIV) 4, the neurons were treated with 1 μM cytosine arabinoside (AraC, Sigma-Aldrich) to prevent a high glial proliferation.

### Treatment of primary hippocampal neuronal cultures

To evaluate the stability and neuronal activity-dependent origin of putative Arc oligomers, cultures were treated with cycloheximide (CHX) to inhibit protein synthesis and tetrodotoxin (TTX) to inhibit neuronal action potential. CHX (50 µg/mL in Milli-Q® water) was applied for 70 min (37 °C), with additional long duration treatments of 3 and 6 hours. Neurons were treated with TTX (2 µM in Milli-Q® water) for 16 h (37 °C), with additional experiments performed with application of 24-72 hours TTX treatment. Neuronal cultures were fixed for imaging after TTX application or following a 30 min washout period with TTX-free medium.

To assess detection of Arc oligomers by PLA, we designed a cell-penetrating, dominant-negative peptide inhibitor of Arc oligomerization, termed OligoOFF. The fluorescein isothiocyanate (FITC) conjugated synthetic peptide has an HIV trans-activator of transcription (TAT) transduction domain and a double glycine linker, followed by 14 amino acids (109 VKRE<u>MHVWREV</u>FYRL 123) from the oligomerization region of Arc’s N-terminal coiled-coil domain [23]. As a control for comparison with the wildtype sequence, we made a TAT peptide with aspartate substitutions in critical residues (M113D and W116D) within the 7-residue oligomerization motif (GenScript Biotech Corporation, Piscataway, NJ) (**Figure 6A**). Cultures were exposed to 1 µM in Milli-Q® water of OligoOFF wildtype or mutant peptide control for 1 hour (37 °C) followed by a 30-min washout to remove peptides that did not enter neurons before fixation and visualization of FITC.

Additionally, on DIV 21, hippocampal neuronal cultures were treated with brain-derived neurotrophic factor (BDNF, Alomone Labs, Jerusalem, Israel, 50 ng/ml in Milli-Q® water, 37 °C) for 2 hours. Moreover, DIV 21 neurons were treated with dihydroxyphenylglycine (DHPG, Tocris/Bio-techne, Minneapolis, MN, 100 µM in Milli-Q® water, 37 °C, for 10 min).

### HEK293FT cell preparation and transfection

The human embryonic kidney 293FT (HEK293FT) cells used in this study were acquired from Thermo Fisher Scientific and were maintained in Dulbecco’s modified Eagle medium (DMEM high-glucose, Sigma-Aldrich) containing 10% FBS and 100 U/mL Penicillin-Streptomycin. When confluent, HEK293FT cells were seeded in coverslips of 12 mm for further performance of the proximity ligation assay (PLA). The cells were transfected with Lipofectamine 2000 Transfection Reagent (Thermo Fisher Scientific) in a concentration of 1 µL/well and 0.5 µg/well of plasmid containing either an empty vector with mTurquoise2 (mTq2) or mTq2-tagged Arc construct. Before fixation of the cells, green fluorescence was checked to confirm transfection of the constructs mentioned above.

### Immunostaining procedures

Prior to the immunostaining protocols for both immunocytochemistry (ICC) and PLA, treated and untreated cells were fixed with 4 % paraformaldehyde, 4% sucrose solution in 0.1 M phosphate buffer, pH 7.4, for 20 min at room temperature.

For the immunocytochemistry protocol, fixed samples were permeabilized with Triton X-100 0.1% in phosphate buffer saline (PBS) for 5 min at room temperature. Following the permeabilization, the samples were washed 3 times with PBS and blocked with 5% goat serum in PBS (blocking solution) for 1 hour at room temperature. After blocking, samples were incubated with anti-drebrin (rabbit polyclonal, Bethyl Laboratories, Inc., Montgomery, TX, 1:1000 dilution) as a neuronal marker, combined with the H11-FLAG tagged nanobody (100 µg/mL) for overnight at 4 °C. On the second day the samples were washed again, three times with PBS and incubated with an anti-FLAG antibody (Mouse monoclonal, Sigma-Aldrich F1804, 1:500 dilution) for 1 hour at 37 °C. Then, the appropriate secondary antibodies: goat anti-mouse IgG (H + L) cross-adsorbed secondary antibody, Alexa Fluor 488 (Thermo Fisher Scientific, dilution 1:500) goat anti-rabbit IgG (H + L) cross-adsorbed secondary antibody, Alexa Fluor 647 (Thermo Fisher Scientific, dilution 1:500), for 1 hour at room temperature, protected from the light. That was followed by another PBS wash and the cover slips were mounted with Prolong Glass Antifade mounting media (Thermo Fisher Scientific) in a microscope glass slide for imaging (Borosilicate glass D 263™ M of hydrolytic class 1, Marienfeld).

For Arc-Arc PLA, both in HEK293FT cells and hippocampal neurons, the Duolink PLA kit (Sigma-Aldrich) was used following the manufacturer’s instructions, with minor changes for the introduction of anti-Arc nanobodies. Briefly, samples were fixed and permeabilized as mentioned above, then blocked with Duolink Blocking Solution for 1 hour at 37 °C. Starting the modified Duolink protocol, coverslips were incubated for overnight at 4 °C with an anti-Arc nanobody H11-FLAG and H11-ALFA or C11-FLAG/C11-ALFA. Concentrations used were 25 µg/mL for H11 and 15 µg/mL for C11 diluted in Duolink Antibody Diluent solution. This was combined with incubation with a primary anti-MAP2 antibody (chicken polyclonal, Encor Biotechnology Inc., Gainesville, dilution 1:3000), used as neuronal marker and to quantify the dendritic area in the image analysis. On the following day, the samples were incubated with anti-ALFA (rabbit polyclonal, NanoTag Biotechnologies GmbH, Göttingen, Germany, N1581, dilution 1:1000) and/or anti-FLAG antibodies (mouse monoclonal, Sigma-Aldrich F1804, dilution 1:500) for 1 hour at 37 °C. Next, the samples were washed and incubated with Duolink In Situ PLA Probes Anti-Rabbit (PLUS) and Anti-Mouse (MINUS) + a secondary antibody Donkey Anti-Chicken IgY (IgG) (H + L) Antibody, CF594A Conjugated (Biotium, Inc., Fremont, CA, dilution 1:500), for 1 hour at 37 °C. Following the probes incubation, the samples were washed and incubated for the ligation step with ligase and a buffer containing oligonucleotides (30 min at 37 °C). The samples were washed after the ligation step and incubated with polymerase and buffer containing fluorophore labelled oligos (Duolink In Situ Detection Reagents FarRed) for the amplification step (100 min at 37 °C). A final washing step was done before mounting the cover slips with Prolong Glass Antifade mounting media in a microscope glass slide for imaging (Borosilicate glass D 263™ M of hydrolytic class 1, Marienfeld). For the positive (PSD95-Stg) and negative (TIP60-Stg), the conventional Duolink manufacturer protocol was used, where the nanobodies were absent.

To determine the concentration used for the nanobody based PLA, a concentration curve (15 to 300 µg/mL) was performed (same protocol as described above, using Duolink PLA Kit) to optimize the use of nanobody H11 pair. The concentration of 25 µg/mL was selected for further testing and validation, considering the best signal/noise ratio observed in the **Supplementary Figure 1**. The concentration of 15 µg/mL was selected for the nanobody C11, due to a higher noise at 25 µg/mL concentration.

### Imaging and image analysis

Except for the images from Arc-Arc PLA in HEK293FT cells, all the other samples (hippocampal neurons) were imaged on AxioImager Z1 microscope (Carl Zeiss AG, Oberkochen, Germany). The HEK293FT cells were imaged on a Leica TCS SP8 confocal microscope (Leica microsystems GmbH, Wetzlar, Germany). The images acquired were all analysed and processed in the software ImageJ (National Institutes of Health, Bethesda, MD).

In both ICC and PLA imaging, random, healthy, and non-overlapping neurons (to distinguish individual dendrites) were selected throughout the coverslip to attain a good representation of each sample. Arc imaging was performed after selection of neurons, thus avoiding selection bias due to Arc immunostaining or PLA. As criteria for selection of neurons for image analysis, the neuronal markers drebrin was used to visualize neuronal spines and dendrites morphology and structure for ICC. For selection of neurons in PLA experiments, MAP2 immunostaining was used to identify neuronal dendrites, as the mouse anti-drebrin antibody cannot be combined with use of anti-mouse PLA probes.

For the Arc ICC analysis, random non-overlapping secondary dendrites from the selected neurons were used. To set the threshold, the background was measured individually in each image and multiplied by a common factor for each experimental replicate to better exclude noise. Puncta counts were normalised by the dendritic length using the formula: (number of puncta x 100 μm)/dendritic length (μm), expressed in Arc puncta/100 μm of dendritic length in the graphs. For all the PLA analysis, MAP2 staining was used to label neurons and select the dendritic area of analysis. First, the neuronal soma was manually removed and then the whole dendritic area was selected using MAP2 staining as reference. We focused our analysis on dendritic branches given long-standing evidence for enrichment of Arc in synaptic and cytoskeletal fractions [43, 44]. The detection threshold was set manually to exclude noise and count PLA signals in the dendrites and peri-dendritic area to include spines. The results were normalized by dendritic area, expressed in number of PLA puncta/dendritic area in μm^2^.

### Statistical Analysis

All the data was processed and analysed using the software GraphPad Prism 10.0.3. The D’Agostino & Pearson test was used to assess data distribution normality for selection of the post-hoc test. Outlier detection was performed using the ROUT method (Q=1%). A minimum of 3 experimental replicates were performed, where a minimum of 20 (ICC) or 45 (PLA) images of individual neurons per group were used for analysis.

One-way ANOVA was used for the analysis of Arc puncta, followed by Dunnett’s multiple comparison test for analysis containing 3 groups. Mann-Whitney or unpaired *t*-test were applied for the comparison between 2 groups, based on normality of the data distribution. For the PLA analysis, for experiments containing 2 groups, Mann-Whitney was applied, and for those containing 3 or more groups, one-way ANOVA followed by Dunnett’s multiple comparison test was applied for normal distribution and Kruskal-Wallis test followed by Dunn’s multiple comparison test for non-parametric tests when suitable.

## Results

### Anti-Arc nanobody H11-FLAG detects endogenous Arc protein via immunocytochemistry

First, we sought to apply H11-FLAG for detection of Arc protein via immunocytochemistry. In untreated DIV 21 hippocampal primary neuronal cultures (**Figure 1A**, **row 1**), sparsely distributed Arc immunoreactive somatodendritic puncta were observed. BDNF treatment significantly increased the density of Arc puncta in dendrites and spines (**Figure 1A**, **2^nd^ row**), Non-specific binding of secondary antibodies was evaluated by omitting H11-FLAG from the protocol in BDNF treated neurons. In the absence of H11-FLAG only negligible background fluorescence was observed (**Figure 1A**, **3^rd^ row**). The quantitative comparison between the groups in presented in **Figure 1C**. Thus, immunocytochemical staining with H11-FLAG detects basal and BDNF-induced Arc as discrete puncta in neuronal dendrites and spines, consistent with enhanced Arc expression detected by antibody-based ICC [7, 45–47].

**Figure 1.**
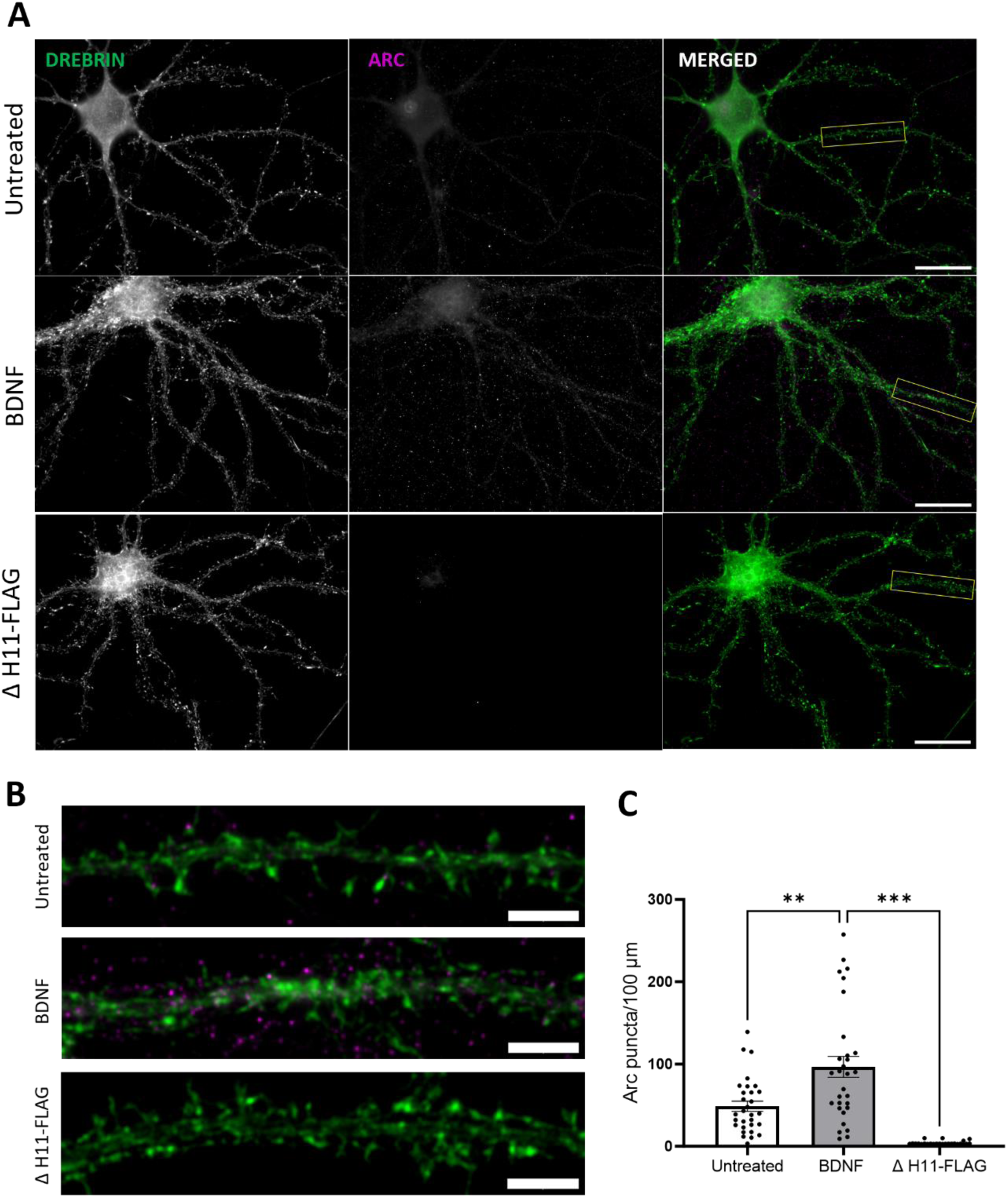
Use of anti-Arc H11-FLAG nanobody for immunocytochemistry in primary hippocampal neuronal cultures. **A** H11-FLAG immunocytochemistry detects an increase in Arc puncta in BDNF treated neuronal cultures compared to unstimulated cultures (untreated), with a low noise signal detected in the absence of H11-FLAG (ΔH11-FLAG). 63x magnification, 25 µm scale bar. **B** Digital zoom of secondary dendrites indicated by a white box in panel A (5 µm scale bar). Drebrin immunocytochemical staining in green and Arc in magenta. **C** Quantification of Arc puncta/100 μm dendritic length ± SEM (30 images/group). One-way-ANOVA followed by Dunnett’s multiple comparisons test; **p≤0.01; ***p≤0.001.

### Nanobody-based PLA detects Arc-Arc complexes in dendrites of unstimulated neuronal cultures

We considered that specific binding of H11 to the Arc N-lobe pocket could be used to develop a PLA method for detection of Arc-Arc interaction complexes, and possible oligomers. For this purpose, H11-FLAG and H11-ALFA were used to target the same epitope in two different Arc molecules. In classical PLA, detection of proximity between two proteins is based on the use of primary antibodies from different species, which are recognized by species-specific secondary antibodies conjugated to DNA oligos. If the distance between the two antibody binding sites is less than 40 nm, oligonucleotide strands can undergo ligation, allowing rolling circle DNA synthesis and detection of the amplicon by fluorescent protein conjugated to complementary oligonucleotide probes. Here, epitope-tagged nanobodies were used to bind Arc, followed by a standard PLA protocol using primary antibodies against the ALFA and FLAG epitopes (**Figure 2**).

**Figure 2.**
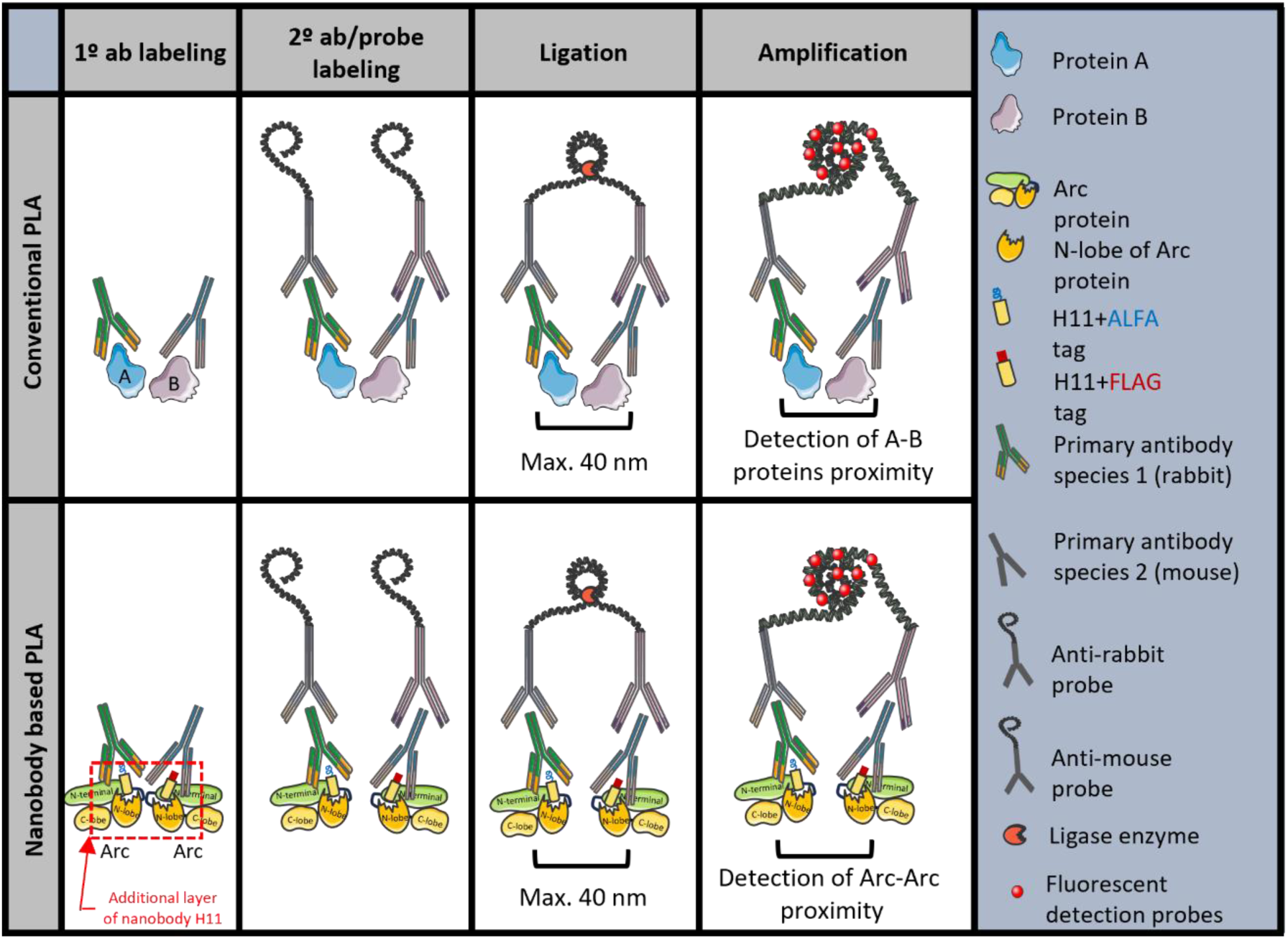
Schematic illustration of conventional and nanobody-based PLA. In conventional PLA, the proteins of interest (A and B) are labelled with a 1° antibody (ab). In nanobody-based PLA for detection of Arc-Arc complexes, we used nanobodies that bind to a single, structurally defined epitope on Arc. Nanobody H11 (depicted here) binds inside the Arc N-lobe ligand binding pocket (dashed red box). The nanobodies are C-terminally fused to ALFA or FLAG tags, which are recognized by rabbit and mouse antibodies, respectively. Subsequent steps for conventional and nanobody-based PLA are the same: Labeling with 2° ab/probes conjugated with oligonucleotides, ligation of oligonucleotides when target proteins are in close proximity (max. 40 nm), addition of DNA polymerase for rolling cycle amplification of ligated DNA, followed by the binding of multiple fluorescent detection probes to create a bright spot.

As positive and negative controls for detection of protein interactions in hippocampal neurons we used standard antibody-based PLA. For a positive control we used the interaction between post-synaptic density protein-95 (PSD-95) and stargazin (Stg) which is an auxiliary subunit of AMPAR-type glutamate receptors. As a mechanism for tethering AMPARs to postsynaptic membranes in dendritic spines, the PSD95-Stg interaction is highly specific and abundant [48, 49]. As a negative control we paired Stg with the nuclear histone acetylase, Tat-interactive protein 60 (TIP60). As expected, a widespread PLA signal was observed along dendrites for PSD95/Stg, whereas almost no signal was detected in the TIP60-Stg PLA (**Figure 3A and 3B**).

**Figure 3.**
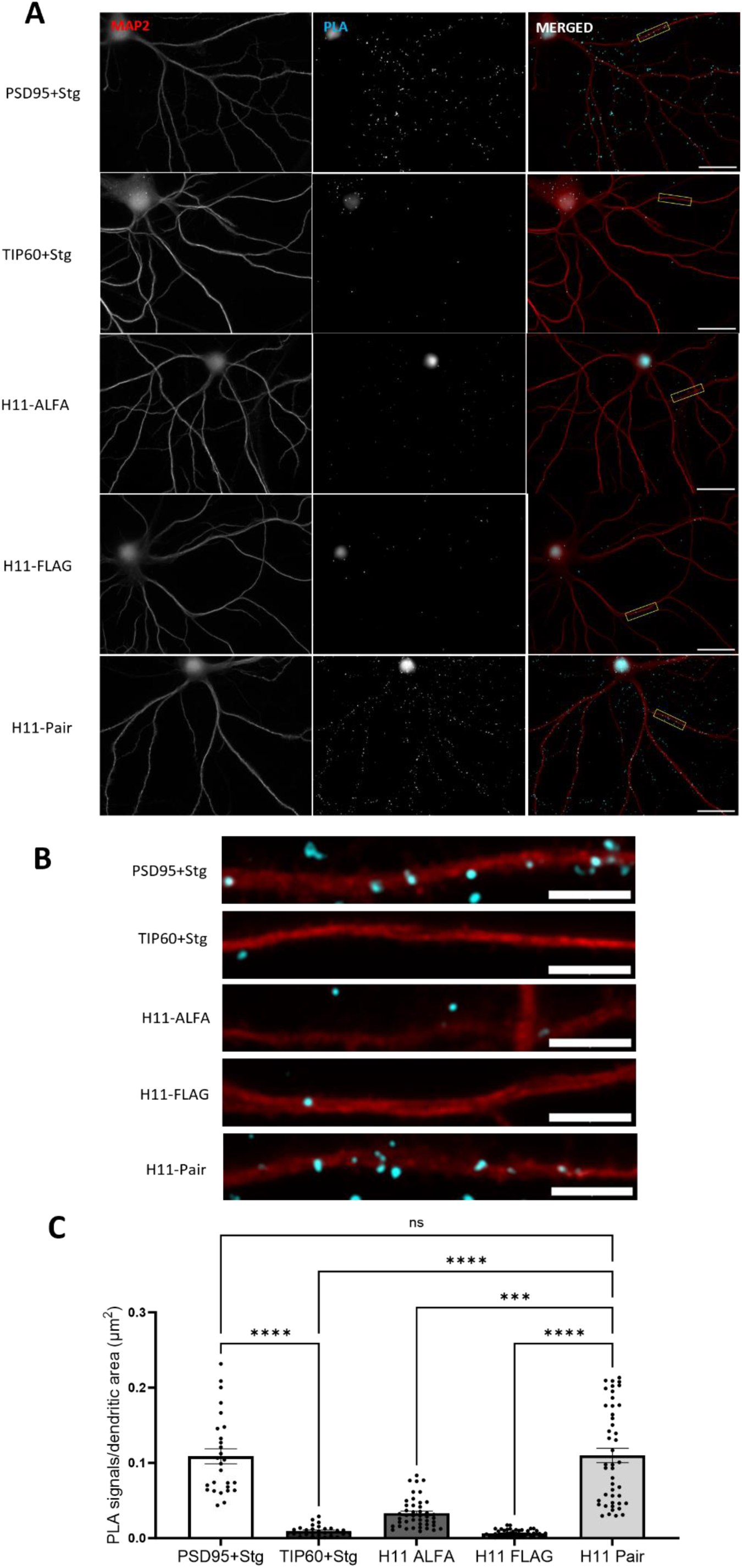
H11 nanobody based PLA detects Arc-Arc complexes in hippocampal neuronal dendrites of unstimulated cultures. **A** Representative image from positive control (PSD95+Stg) and negative control (TIP60+Stg) antibody-based PLA; nanobody PLA using only ALFA-H11 or FLAG-H11 FLAG (rows 3 and 4, respectively), and nanobody PLA using the ALFA-H11/FLAG-H11 pair. Scale bar is 25 µm. **B** Digital zoom of secondary dendrites indicated by a white box in panel A (5 µm scale bar). **C** Quantification of PLA signals/dendritic area (μm^2^) ± SEM (30-45 images/group). One-way-ANOVA followed by Dunnett’s multiple comparisons test showing ***p≤0.001 and ****p≤0.0001.

In the H11 nanobody-based PLA, we observed a widespread signal in dendrites like that observed in PSD95-Stg positive control, suggesting ubiquitous detection of Arc-Arc complexes. When either H11-ALFA or H11-FLAG was omitted from the protocol, only sparse PLA signals were detected, similar to the TIP60-Stg negative control. To quantify the difference observed in the images, the neuronal marker, MAP2 was used to randomly select neurons. PLA puncta were counted using ImageJ software and normalized by dendritic area determined using MAP2 staining, focusing only on the dendritic and synaptic signals, expressed as PLA puncta/dendritic area (μm^2^). PLA signal density in the H11 ALFA/FLAG pair protocol was equivalent to that of the PSD95-Stg positive control and significantly higher than the TIP60-Stg negative control and the H11-ALFA or H11-FLAG alone controls. (**Figure 3C**; One-way-ANOVA followed by Dunnett’s multiple comparisons test showing p>0.999). These results show that Arc-Arc interactions detected by H11 PLA are equivalent in abundance to PSD-Stg interactions in hippocampal neuronal cultures.

To further validate H11 specific labelling of Arc-Arc interaction we used HEK293FT cell line (**Supplementary Figure 2**). HEK293FT cells do not express endogenous Arc, but ectopically expressed mTq2-tagged Arc in these cells is detected as monomers and oligomers by in situ protein crosslinking analysis [23]. Using the H11 nanobody pair, cells transfected with mTq2-tagged Arc exhibited clear and abundant PLA signals, while non-transfected cells or cells transfected with empty vector (mTq2) show sparse signals, equivalent to background noise. Similarly, in cells expressing mTq2-tagged Arc, only sparse signal was detected when either of the epitope-tagged H11 nanobodies was omitted.

### Basal Arc-Arc complexes are independent of recent neuronal activity and protein synthesis

Arc is known for its rapid activity-dependent transcription, translation, and rapid protein degradation, with protein half-life estimates of 0.5 to 1 hour [11–13, 50]. In unstimulated hippocampal neuronal cultures, expression of Arc mRNA and other immediate early genes occurs in a subfraction of neurons [51–53]. The ubiquitous Arc-Arc PLA signal found in unstimulated neuronal cultures is therefore surprising and suggests the existence of constitutive Arc complexes. To assess the origin and stability of Arc complexes, tetrodotoxin (TTX) was used to block action potentials (**Figure 4**), and cycloheximide (CHX) was used to inhibit protein synthesis (**Figure 5**).

**Figure 4.**
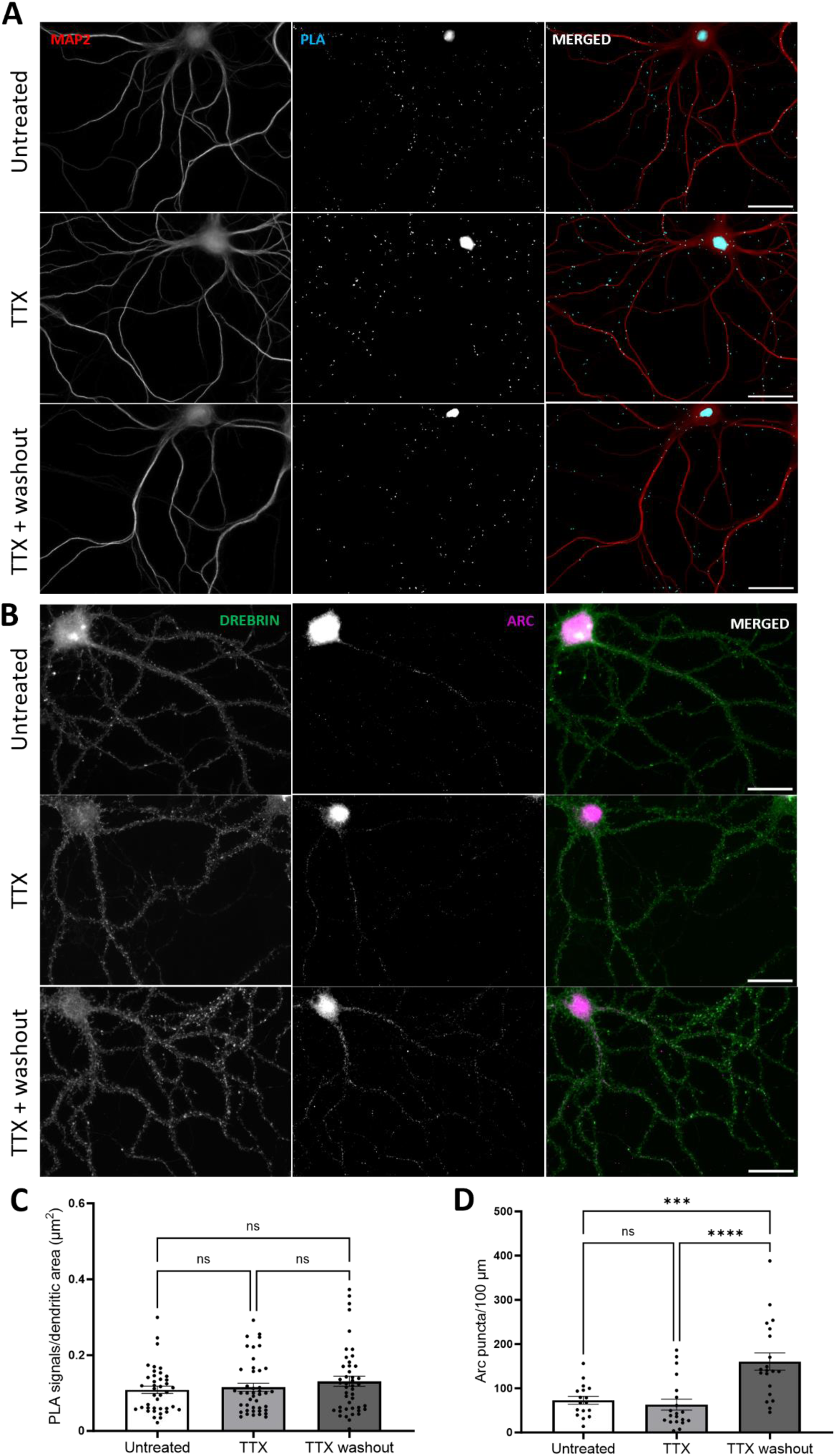
Inhibition of neuronal firing by TTX does not affect basal Arc-Arc complexes. **A** Representative Arc-Arc H11 PLA images from hippocampal neuronal cultures treated with TTX for 16 h, TTX followed by 30 min washout, or no treatment. **B** Representative immunocytochemical staining using anti-Arc H11-FLAG nanobody. **C** Quantification of the comparison between treated (TTX and TTX + washout, 2^nd^ and 3^rd^ rows, respectively) groups and untreated groups, expressed in PLA signals/ dendritic area (μm2) ± SEM (45 images/group). Kruskal-Wallis test followed by Dunn’s multiple comparisons test showing p≥0.999 for both groups. **D** Quantification of the ICC comparing treated groups (TTX and TTX + washout, 2^nd^ and 3^rd^ rows, respectively) and untreated groups, expressed in Arc puncta/100 μm dendritic length ± SEM (20 images/group). One-way-ANOVA followed by Dunnett’s multiple comparisons test showing ***p≤0.001 and **** p≤0.0001. Scale bar of 25 µm.

**Figure 5.**
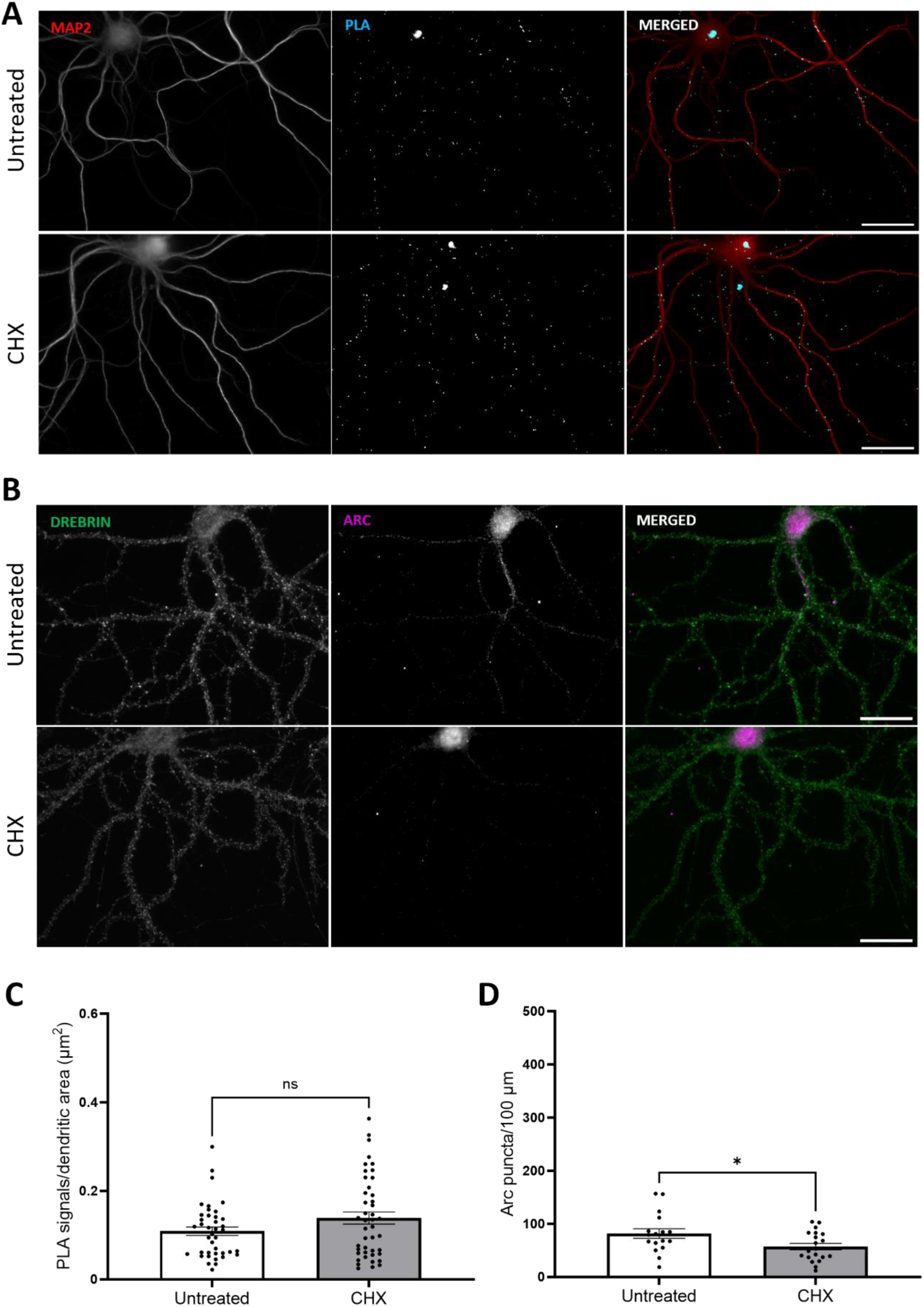
Inhibition of protein synthesis does not affect basal Arc-Arc complexes. **A** Representative image of treated and untreated groups for PLA detection. **B** Representative images of treated and untreated groups for ICC. **C** Quantification of the comparison between treated and untreated groups, expressed in PLA signals/ dendritic area (μm2) ± SEM (45 images/group). Mann Whitney test showing p=0.21. **D** Quantification of the ICC comparison between treated and untreated groups, expressed in Arc puncta/100 μm dendritic length ± SEM (20 images/group). Unpaired T test showing *p≤0.05. Scale bar of 25 µm.

Silencing of neuronal firing with TTX is known to inhibit Arc transcription, while washout of TTX results in rebound of neuronal activity and enhanced transcription [51]. We found that TTX treatment (2 µM) for 16 h did not affect the abundance of Arc-Arc PLA signal (**Figure 5C**). Washout for 30 min post-TTX also did not alter PLA signal (**Figure 4A and 4C**), although ICC analysis (**Figure 4B and 4D**) with H11 nanobody demonstrated a robust increase in Arc puncta density in hippocampal neuronal dendrites. Longer TTX treatments for 24, 48, and 72 hours also had no effect on the Arc-Arc PLA signal abundance (**Supplementary Figure 3**).

CHX 50 µg/mL treatment for 70 min also did not alter Arc-Arc PLA signal abundance in neuronal dendrites (**Figure 5C**), even though CHX significantly reduced Arc expression (**Figure 5D**) as detected by ICC staining. Even extended periods of CHX treatment of 3 and 6 hours did not significantly alter the PLA (**Supplementary Figure 4**). Thus, inhibition of recent neuronal activity and protein synthesis affects expression of Arc but does not alter the abundance of the Arc/Arc PLA signal, suggesting the presence of stable Arc complexes in neuronal dendrites.

### Arc-Arc PLA signal is inhibited by a cell-penetrating peptide inhibitor of Arc oligomerization

Next, we asked whether stable Arc complexes reflect the detection of oligomeric Arc. The Arc N-terminal coiled-coil domain plays a critical role in Arc self-association and oligomerization. Mutation of the oligomerization motif in coil-2 inhibits the assembly of dimers into higher-order oligomers [23]. We therefore designed a cell-penetrating TAT peptide (OligoOFF) as a dominant-negative inhibitor of Arc oligomerization. OligoOFF has 14 residues from the coil-2 oligomerization region harboring the 7-residue oligomerization motif (a.a. 113-119, MHVWREV) (**Figure 6A**). M113 and W116 are critical residues in the Arc-Arc atomic interface, and aspartate mutation of these residues inhibits oligomerization [23]. As a control TAT peptide, we therefore substituted M113 and W116 with aspartate.

**Figure 6.**
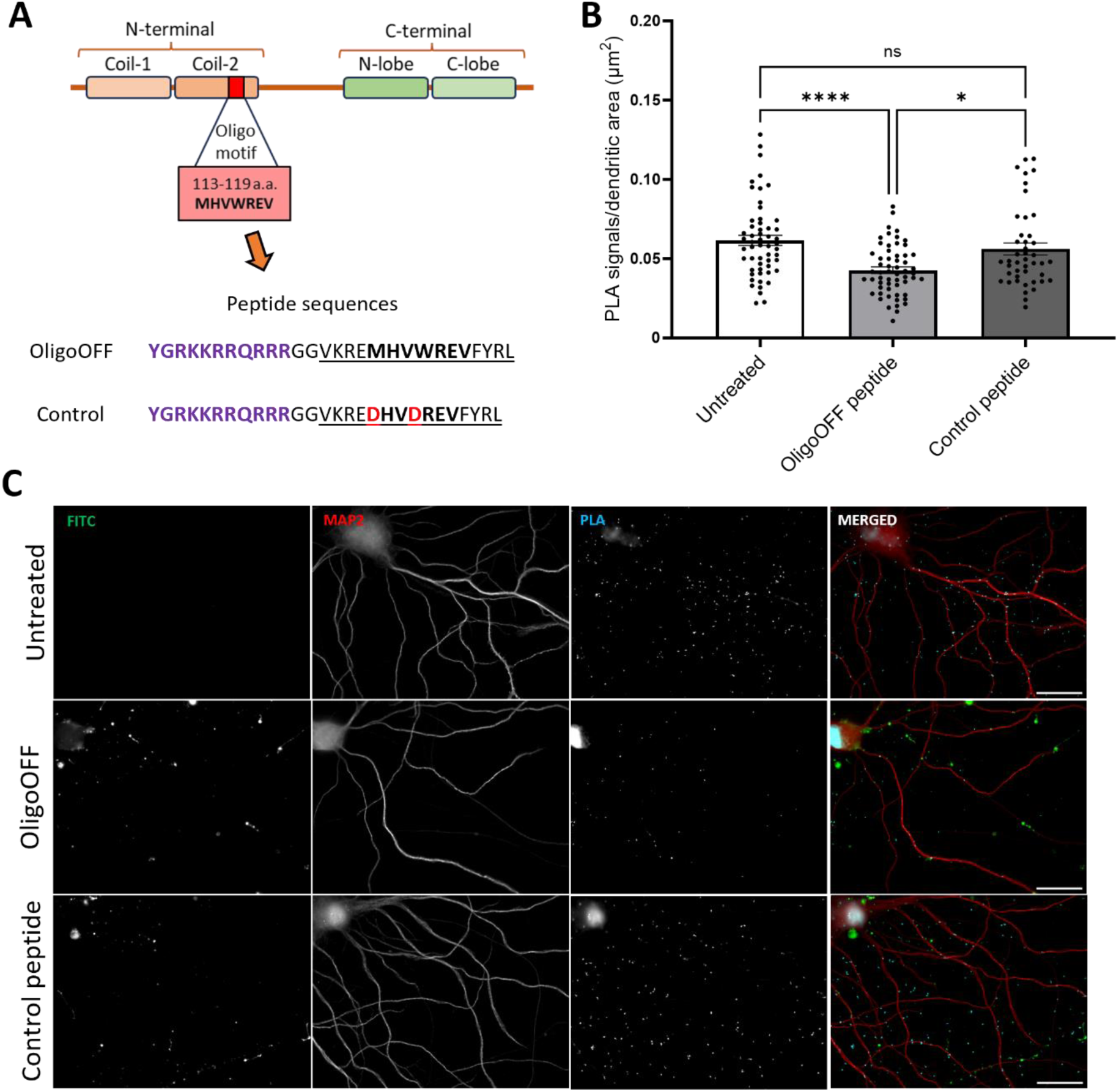
Arc-Arc complexes detected by H11 PLA are inhibited by a cell-penetrating peptide inhibitor of Arc oligomerization. **A** Cell penetrating peptide dominant-negative inhibitor of Arc oligomerization motif. Top panel. Arc domain structure illustrating position of 7-residue oligomerization motif. Bottom panel. Amino acid sequence of cell-penetrating peptides. TAT sequence in purple, double glycine linker, followed by wildtype sequence of oligomerization peptide (underlined) and oligomerization motif (in bold). In control peptide, critical residues M113/Trp116 were replaced with aspartate (in red). **B** Quantification of comparison between OligoOFF group and untreated or control peptide group, expressed in PLA signals/ dendritic area (μm^2^) ± SEM (45-60 images/group) (Kruskal-Wallis test followed by Dunn’s multiple comparisons test showing ****p≤0.0001 and *p≤0.05) detection. Scale bar of 25 µm. **C** Representative images of untreated and groups treated OligoOFF and control peptide, respectively.

PLA detection was performed with H11 after incubation of cultures with fluorescent FITC-conjugated TAT peptide for 1 hour. Both OligoOFF and control peptide penetrated the neuronal cell body and dendrites, as shown by FITC fluorescence in **Figure 6C**. Application of OligoOFF strongly reduced the PLA signal relative to untreated neurons and cultures treated with control peptide (**Figure 6B**). In addition, there was no difference in PLA signal between the control peptide treated cultures and untreated cultures, demonstrating the importance of the intact oligomerization motif for inhibition of Arc-Arc interaction (**Figure 6B**). These results suggest that H11 PLA signal reflects the detection of constitutive, stably expressed oligomeric Arc.

### Nanobody PLA detects Arc-Arc complexes induced by BDNF and DHPG

Next, we asked whether Arc complexes are regulated by stimuli that are known to induce Arc expression. As BDNF enhances Arc expression in hippocampal neuronal dendrites (**Figure 1**), we expected to observe enhanced Arc-Arc PLA following BDNF treatment. Surprisingly, BDNF treatment significantly reduced signal abundance detected by H11 PLA (**Figure 7C**). Although H11 detects constitutive Arc complexes, it might not detect all oligomeric forms. We therefore employed another anti-Arc nanobody, clone C11, which binds to an exposed surface of the capsid domain C-lobe [37]. First, we show that PLA performed with C11-ALFA and C11-FLAG detects constitutive Arc-Arc complexes in hippocampal neuronal dendrites as observed with H11 PLA (**Supplementary Figure 5**). Again, treatment with TTX and CHX did not affect the constitutive Arc-Arc PLA signal density (**Supplementary Figure 6**). However, in contrast to H11, the C11-based PLA detected enhanced Arc-Arc interaction in BDNF treatment neurons (**Figure 7**).

**Figure 7.**
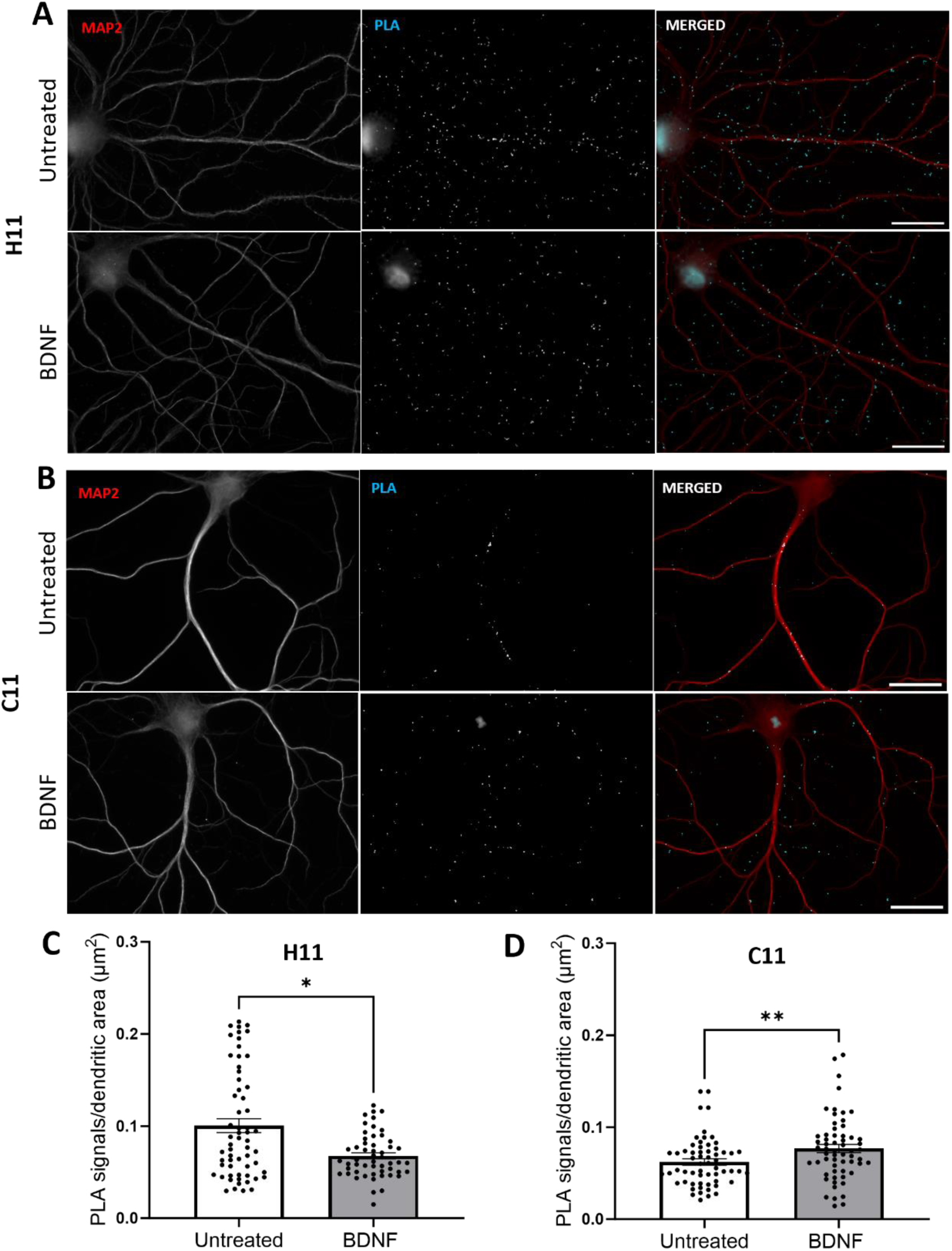
BDNF increases Arc-Arc PLA signal detected by the C-lobe binding nanobody C11 but not by H11. **A** Representative image of untreated and BDNF-treated groups detected by H11 pair. **B** Representative images of untreated and BDNF-treated groups detected by H11 pair. **C** Quantification of the comparison between control and BDNF groups, expressed in PLA signals/ dendritic area (μm^2^) ± SEM (60 images/group) detected by H11 nanobody (Mann Whitney test showing *p<0.05). **D** Quantification of the comparison between control and BDNF groups, expressed in PLA signals/dendritic area (μm^2^) ± SEM (60 images/group) detected by C11 nanobody (Mann Whitney test showing **p<0.01). Scale bar of 25 µm.

We considered that epitopes for H11 (N-lobe ligand pocket) and C11 (C-lobe surface) are sensitive to Arc oligomeric state (**Figure 8A**). Previous work using *in situ* protein crosslinking to trap Arc-Arc interactions, showed that BDNF treatment of hippocampal neurons dramatically enhances expression of Arc monomers and dimers, with no evidence of tetramers [32]. The present results could be explained by preferential detection of dimer by C11 and higher-order oligomer by H11. To test this hypothesis, we repeated the OligoOFF peptide experiment in non-stimulated neuronal cultures using C11 for PLA (**Figure 8B**). In contrast to H11, the C11 PLA showed no difference in signal density between OligoOFF treated and untreated cultures (**Figure 8C**). Notably, the oligomerization motif mediates dimer assembly into higher-order oligomers, but not dimer formation itself [23]. Therefore, the results with C11 are consistent with detection of BDNF induced dimers and persistence of dimers following disruption of constitutive Arc-Arc complexes by OligoOFF.

**Figure 8.**
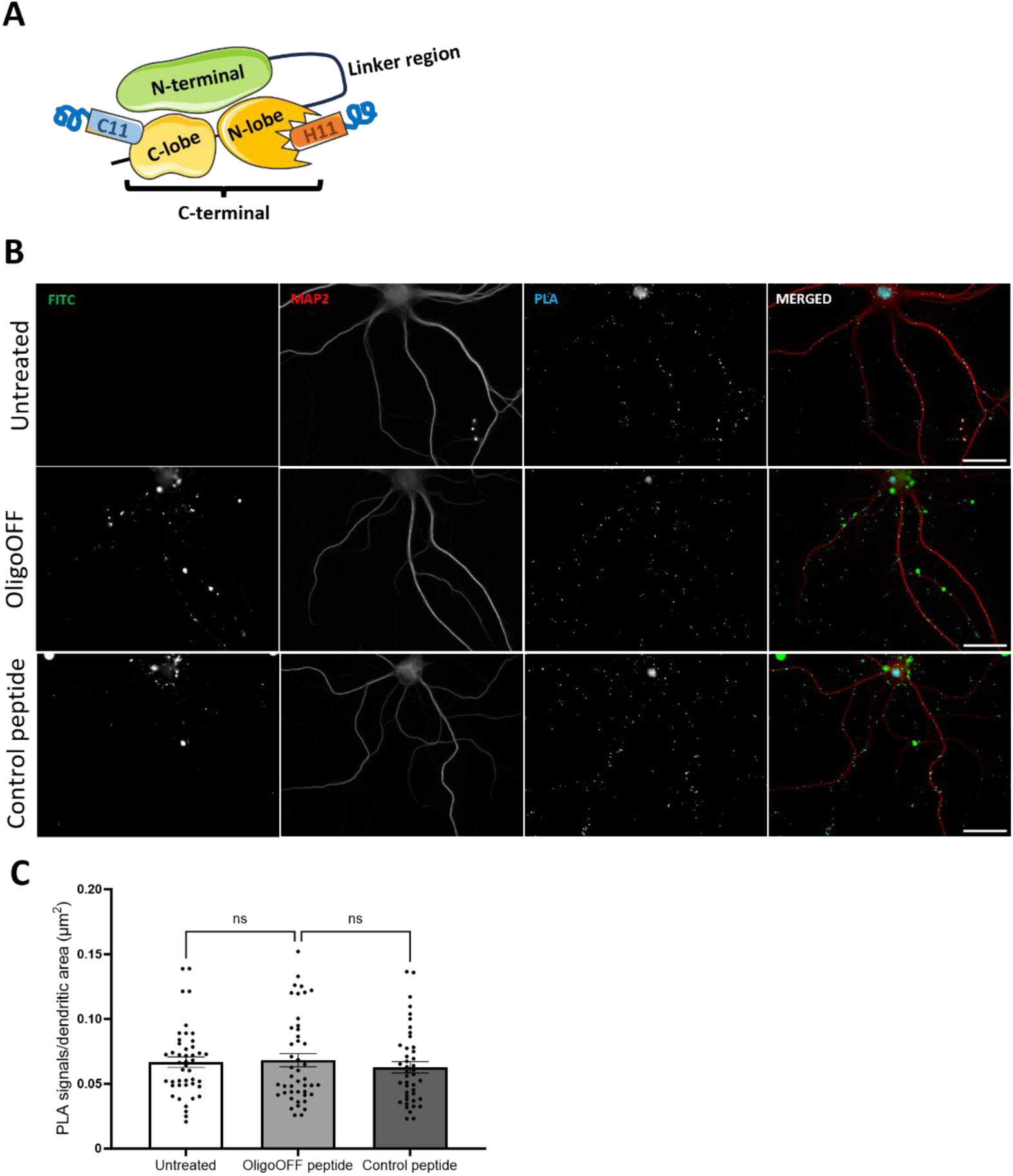
Arc-Arc complexes detected by C11 are not inhibited by OligoOFF-peptide. **A** Cartoon illustrating anti-Arc nanobody H11-ALFA binding to the N-lobe ligand pocket and C11-ALFA binding to the C-lobe surface of the Arc’s C-terminal capsid domain. C11-ALFA and C11-FLAG were used for the PLA data shown here. **B** Representative image of untreated and groups treated OligoOFF and control peptide. **C** Quantification of comparison between OligoOFF group and untreated or control peptide group, expressed in PLA signals/ dendritic area (μm^2^) ± SEM (45-60 images/group), for C11 detection (Kruskal-Wallis test followed by Dunn’s multiple comparisons test showing no significance in the comparisons). Scale bar of 25 µm.

To compare with BDNF, hippocampal neuronal cultures were treated with the metabotropic glutamate receptors agonist DPHG. DHPG induced Arc synthesis is critical for synaptic depression, and this mechanism is attenuated by mutations that prevent Arc higher-order assembly from tetramers to 32-mers [8, 22, 26]. We investigated whether DHPG stimulation would induce changes in Arc-Arc PLA detected via H11 or C11. (**Figure 9A** and **9B**, respectively). Quantification of PLA signals for nanobody H11 demonstrated a significant increase in signal abundance in the DHPG treatment group relative to untreated control (**Figure 9C**). In contrast, PLA signals for nanobody C11 did not differ between DHPG-treated and untreated control (**Figure 9D**). These results again show differential detection of Arc-Arc complexes H11 and C11 nanobodies, consistent with preferential detection of higher-order oligomers by H11.

**Figure 9.**
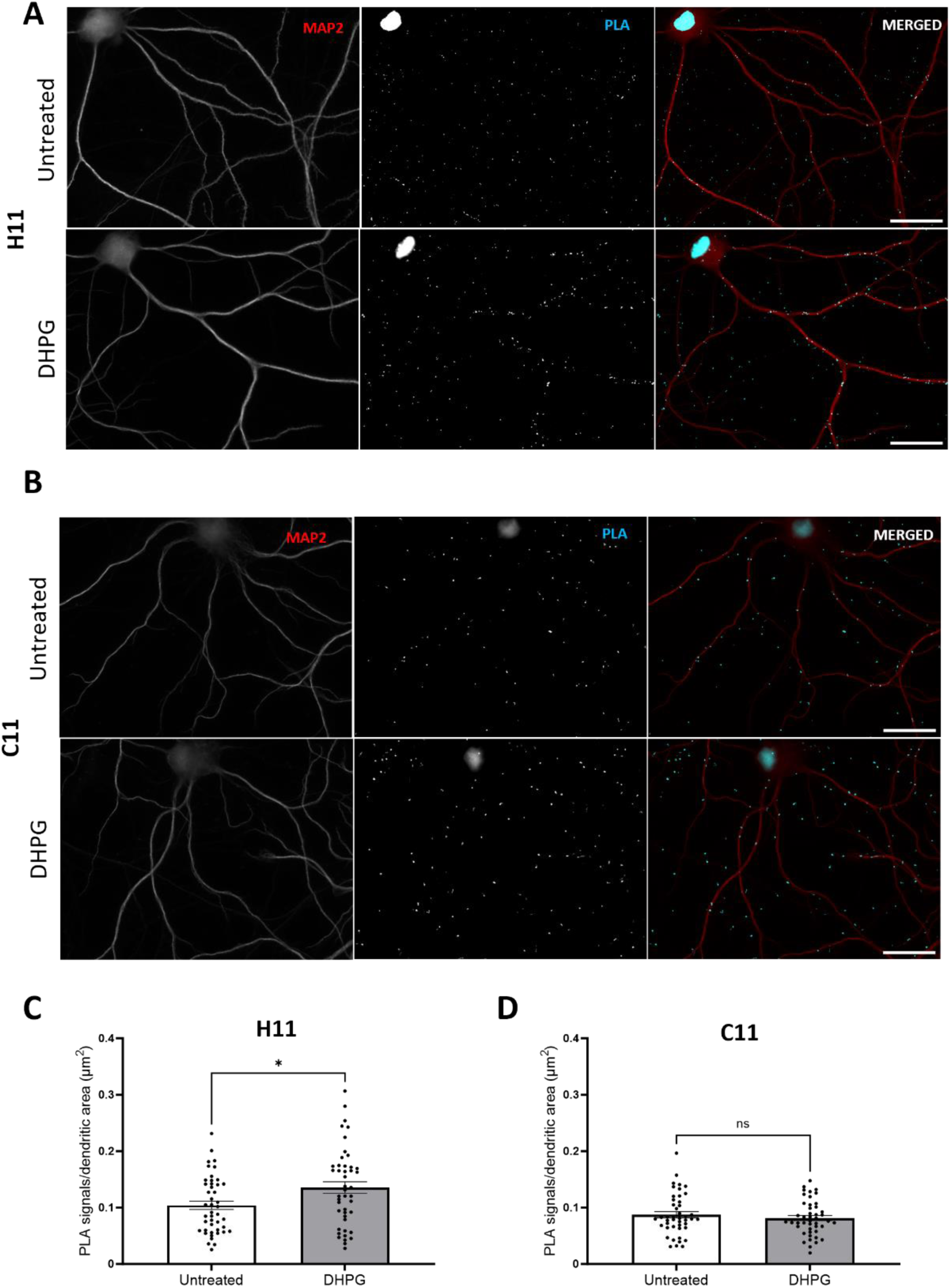
DHPG increases Arc-Arc PLA signal detected by H11 but not C11 nanobody. **A** Representative image of untreated and DHPG-treated groups detected by H11 pair. **B** Representative images of untreated and DHPG-treated groups detected by C11 pair. **C** Quantification of the comparison between control and DHPG groups, expressed in PLA signals/ dendritic area (μm^2^) ± SEM (45 images/group) detected by H11 nanobody (Unpaired T test showing *p<0.05). **D** Quantification of the comparison between control and DHPG groups, expressed in PLA signals/ dendritic area (μm^2^) ± SEM (45 images/group) detected by C11 nanobody (Unpaired Student’s T-test showing p=0.3906). Scale bar of 25 µm.

## Discussion

The novel nanobody-based PLA provides a method for detection, localization, and quantification of native Arc-Arc complexes. Surprisingly, PLA revealed widespread Arc-Arc complexes in neuronal dendrites under basal conditions in unstimulated hippocampal neuronal cultures. The pool of Arc detected by PLA is not reliably detected by ICC staining, indicating the higher sensitivity of the PLA method [39, 40, 54]. The Arc-Arc PLA signal density and distribution is similar to PLA for the Stg-PSD95 interaction in dendritic spines. The basal Arc-Arc PLA signal is also unaffected by inhibition of neuronal activity (TTX treatment for up to 72 hours) and inhibition of protein synthesis (CHX treatment for up to 6 hours), demonstrating constitutive expression of stable Arc-Arc complexes in dendrites or dendritic spines. These complexes can be attributed to oligomeric Arc based on the strong inhibition of H11 PLA signal by OligoOFF Tat peptide, but not control peptide in which two residues critical for Arc-Arc interaction (methionine 113 and tryptophan 116) were changed to aspartate.

In principle, Arc-Arc could reflect binding to partner proteins within crowded environments such as the PSD. Localization of Arc to postsynaptic sites depends on PSD95 and it is estimated that 5% of biochemically identified PSD95 supercomplexes contain Arc and 58% of Arc supercomplexes contain PSD95 [55]. Arc binds through its N-lobe with several PSD proteins including NMDA receptor subunit 2B, stargazin (TARPγ2) and guanylate kinase anchoring protein (GKAP) [19, 56]. In addition, Arc purifies with the crude F-actin fraction [43], and interacts with the actin-binding protein, drebrin A, within the spine core [44]. Under basal conditions, in non-stimulated hippocampal dentate gyrus, subcellular fractionation and co-immunoprecipitation with immunoblot analysis shows that Arc is highly enriched in the synaptic, cytoskeletal fraction in complex with drebrin A [44]. It is therefore possible that constitutive Arc-Arc PLA reflects oligomeric Arc in complex with partner proteins in the spine core and PSD.

Arc oligomeric state has emerged as a determinant of Arc function. However, current functional data in neuronal cultures, brain slices, and live rodents, is based on mutations that impair oligomerization of purified Arc protein, while regulation of endogenous Arc oligomers is largely unknown. The present study provides evidence for stimulus-evoked increases in Arc-Arc complexes in response to both BDNF and DHPG. Moreover, differential detection of stimulus-induced complexes by H11 and C11 indicate recognition of distinct oligomeric forms or altered access of nanobodies to binding sites. We show that BDNF treatment induces an increase in C11 but not H11 PLA signal abundance, whereas DHPG induced Arc-Arc complexes are detected by H11 but not C11 PLA. Using in situ protein cross-linking of hippocampal cultures to capture weak non-covalent interactions, Mergiya and collaborators (2023) found several fold increases in Arc monomer and dimer following BDNF treatment, with no apparent increase in tetramer or other larger species [32]. BDNF induces Arc-dependent LTP [6, 7], whereas DHPG induces Arc synthesis-dependent LTD. Zhang and colleagues (2019) showed reduction of DHPG-induced LTD in mice harbouring mutations that impair oligomerization from tetramer to 32mers in the recombinant Arc protein. Together these data are consistent with a working hypothesis presented in Figure 10, in which C11 PLA detects Arc dimer and H11 higher-order oligomers. Basal Arc complexes are detected in dendrites at equal density by both H11 and C11 PLA, but only H11 PLA is disrupted by OligoOFF peptide. OligoOFF targets the N-terminal motif which is critical for dimer interactions towards higher-order oligomers and capsids but does not mediate dimer formation itself [23]. Accordingly, the differential effect of OligoOFF also supports preferential detection of dimers by C11 and higher-order oligomers by H11 PLA.

**Figure 10.**
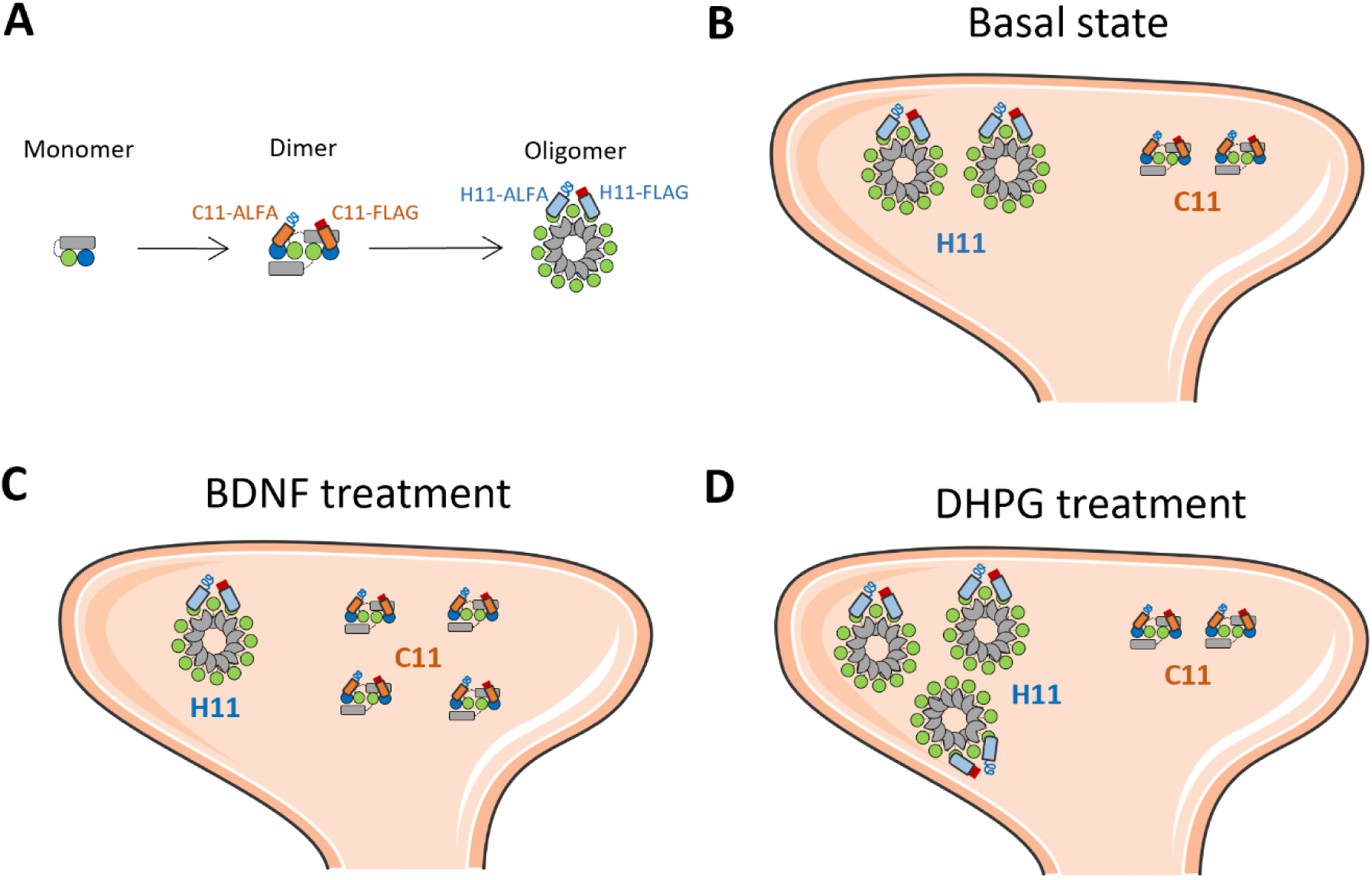
Working hypothesis of constitutive and stimulus-induced Arc oligomers detected by H11 and C11 PLA. **A** Schematic cartoon of Arc protein domains in monomer, dimer, and higher-order oligomer. Arc NTD (gray rectangle), CTD capsid domain with N-lobe (green circle) and C-lobe (blue circle). The dimer is the building block for higher-order assembly. The figure depicts a putative domain-swapped stable dimer and a higher-order oligomer. We propose that H11 PLA preferentially detects higher-order oligomer with multiple, accessible N-lobe ligand binding sites, while C11 PLA preferentially detects dimer. **B** Basal state showing constitute expression of dimer and high-order oligomer in a dendritic spine. Both H11 and C11 PLA detect Arc-Arc complexes in the neuronal basal state. **C** BDNF treatment increases Arc-Arc C11 PLA signal while decreasing H11 signal, consistent with enhanced *de novo* synthesis of dimer and either disassembly of higher-order oligomer or blocked access of H11 to the N-lobe pocket. **D** DHPG treatment increases H11, but not C11, PLA signal, consistent with enhanced selective formation of higher-order oligomers.

H11 and C11 binds to Arc with low nanomolar affinity and crystal structures of the nanobodies in complex with the capsid domain have been solved. The H11 complementarity determining region 3 binds inside the ligand pocket through a conserved N-lobe binding motif. C11 binds to an exposed surface of the C-lobe formed by helix-6 and -7 which includes conserved sites of C-lobe/C-lobe inter-subunit binding important for retroviral capsid formation [37]. Clearly, the regulation of H11 and C11 PLA signals indicates major difference in the availability of epitopes for binding in native Arc-Arc complexes. The H11 PLA results indicate enhanced access to N-lobe (green circles) hydrophobic pockets in higher-order oligomers (**Figure 10**) that is lost or dispersed upon disruption by OligoOFF. In turn, the C11 results suggests that C-lobe (blue circles) surfaces are not exposed for PLA interaction in the larger oligomers nor are they created after disruption of complexes by OligoOFF. Clearly, complex interaction between exposure of vacant binding sites and occlusion of sites by endogenous ligands may occur. For instance, the observed reduction in H11 PLA following BDNF treatment could reflect enhanced binding of constitutive oligomers to N-lobe binding partners at synapses or release of capsids from the neuron, in parallel with the observed enhanced synthesis of Arc and increased abundance of C11 PLA signal.

## Conclusion

The present study introduces nanobody-PLA as a method for detecting native Arc-Arc complexes, including oligomers, in neuronal cultures in situ. Using nanobodies that recognize structurally defined epitopes in the Arc capsid domain, we identified widespread constitutive Arc-Arc oligomers in neuronal dendrites that are stably expressed and not dependent on recent neuronal firing activity or protein synthesis. We also uncover regulation of Arc-Arc complexes specific to BDNF and DPHG stimuli. This work thus places focus on the role of constitutive oligomeric Arc and dynamic regulation oligomers in the context of neuronal signalling and plasticity.

## Supplementary Information

### Ethics approval

All animal procedures and handling were performed accordingly with the Norwegian National Research Ethics Committee in compliance with EU Directive 2010/63/EU, ARRIVE guidelines.

### Consent to participate

No human subjects nor samples were used in the present study.

### Consent for publication

No human subjects nor samples were used in the present study.

### Availability of data and materials

All data used in this study are available from the corresponding author upon reasonable request.

### Competing interests

All authors declare that there were no known competing financial or personal interests that could have appeared to influence the work reported in the present study.

## Funding

This was work was funded by grants from the Research Council of Norway (249951) and the Trond Mohn Foundation (grant TMS2021TMT04) to CRB. RB was recipient of a PhD fellowship from the Faculty of Medicine, University of Bergen. FP was supported by a postdoctoral fellowship through Norway-Poland EAA grant to CRB. HF was supported by a scholarship from The Medical Student Research Programme at The Faculty of Medicine, University of Bergen.

## Author contributions

Rodolfo Baldinotti, Francois Pauzin, and Clive Bramham designed and supervised the study. Rodolfo Baldinotti developed the nanobody-based PLA and performed the PLA and ICC work with nanobodies H11 and C11. Hauk Fevang performed experiments and data analysis for C11 nanobody PLA for figures 7, 8 and supplementary figure 5. Rodolfo Baldinotti and Francois Pauzin performed image data analysis and statistics. Rodolfo Baldinotti prepared the figures. Rodolfo Baldinotti prepared the hippocampal neuronal cultures, with contribution of cryopreserved neurons from Yuta Ishizuka. Rodolfo Baldinotti and Clive Bramham wrote the manuscript, with contributions from all authors. All authors have read and approved the final manuscript.

## Acknowledgments.

The authors thank Dr. Hongyu Zhang for providing TAT-peptides for this study.

## Abbreviations

AMPAR: α-amino-3-hydroxy-5-methyl-4-isoxazolepropionic acid receptors
ANOVA: analysis of variance
AraC: cytosine arabinoside
Arg3.1: activity-regulated gene 3.1
Arc: activity-regulated cytoskeleton-associated protein
BDNF: brain-derived neurotrophic factor
CHX: cycloheximide
CDR3: complementarity determining region 3
CA: C-terminal capsid domain
DHPG: dihydroxyphenylglycine
DIV: day in vitro
DMEM: Dulbecco’s modified Eagle medium
FBS: fetal bovine serum
Gag: group-specific antigen
HBSS: Hank’s balanced salt solution
HEK: human embryonic kidney cells
ICC: immunocytochemistry
LTD: long-term depression
LTP: long-term potentiation
MAP2: microtubule-associated protein 2
MEM: minimum essential medium
PBS: phosphate buffer saline
PLA: proximity ligation assay
PLL: poly-l-lysine
PSD: post-synaptic density
Stg: stargazin
TAT: trans-activator of transcription
TIP60: Tat-interactive protein 60
TTX: tetrodotoxin

## Notes

### Competing Interest Statement

The authors have declared no competing interest.

