## Supplemental Figures for "A nanobody-based proximity ligation assay detects constitutive and stimulus-regulated native Arc/Arg3.1 oligomers in hippocampal neuronal dendrites"

### Supplementary Figures

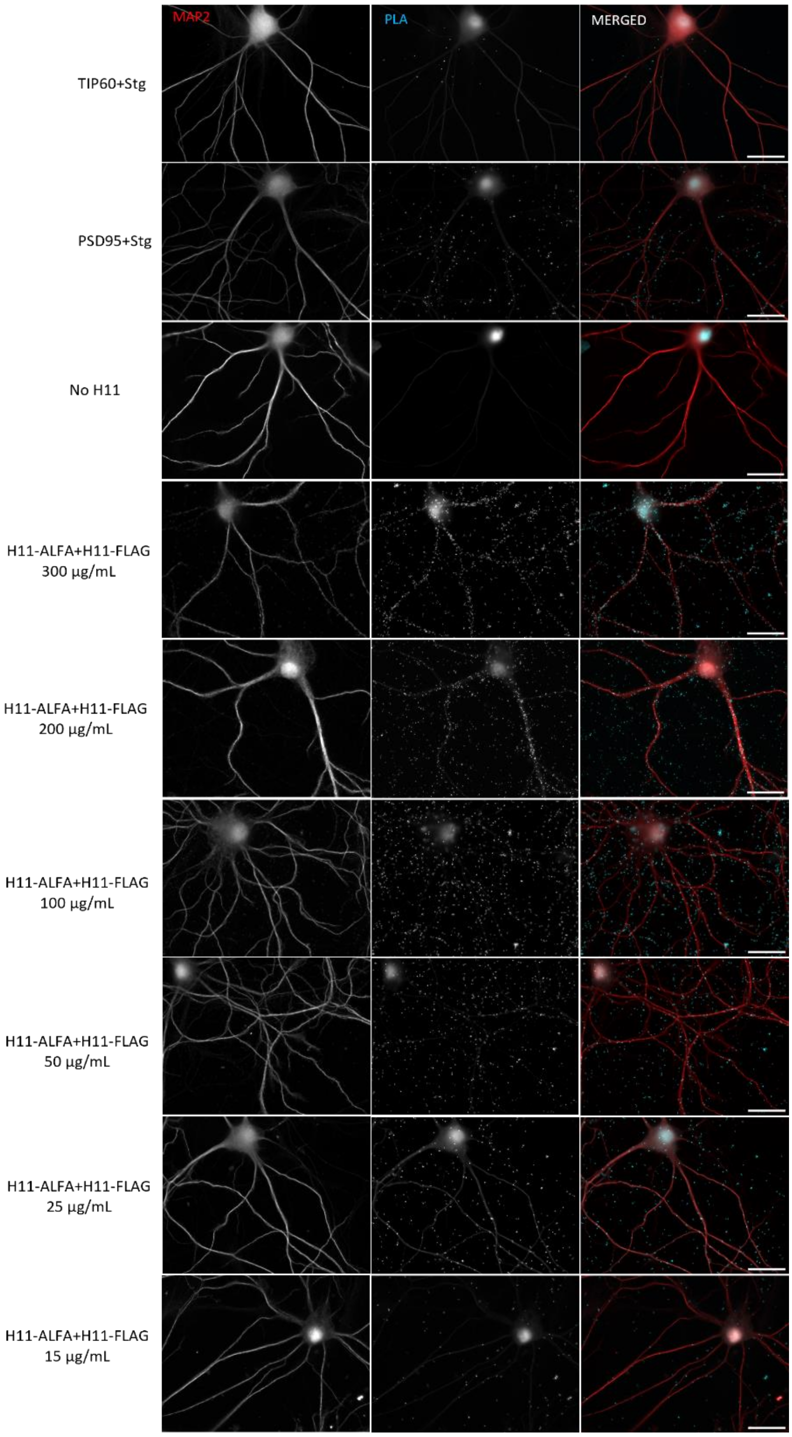


**Supplementary Figure 1. Concentration curve optimization for H11 nanobody-based Arc-Arc PLA validation in hippocampal neuronal primary culture (DIV21).** 1st to the 3rd rows contain 3 different controls groups: (1) positive control (known protein-protein interaction, PSD95+Stg), (2) negative control (TIP60+Stg) and (3) no nanobody H11. From 4th to 9th rows are displayed 6 different concentrations of the nanobody H11-ALFA/FLAG pair (decreasing from 300 to 15 µg/mL, respectively). Scale bar of 25 µm.

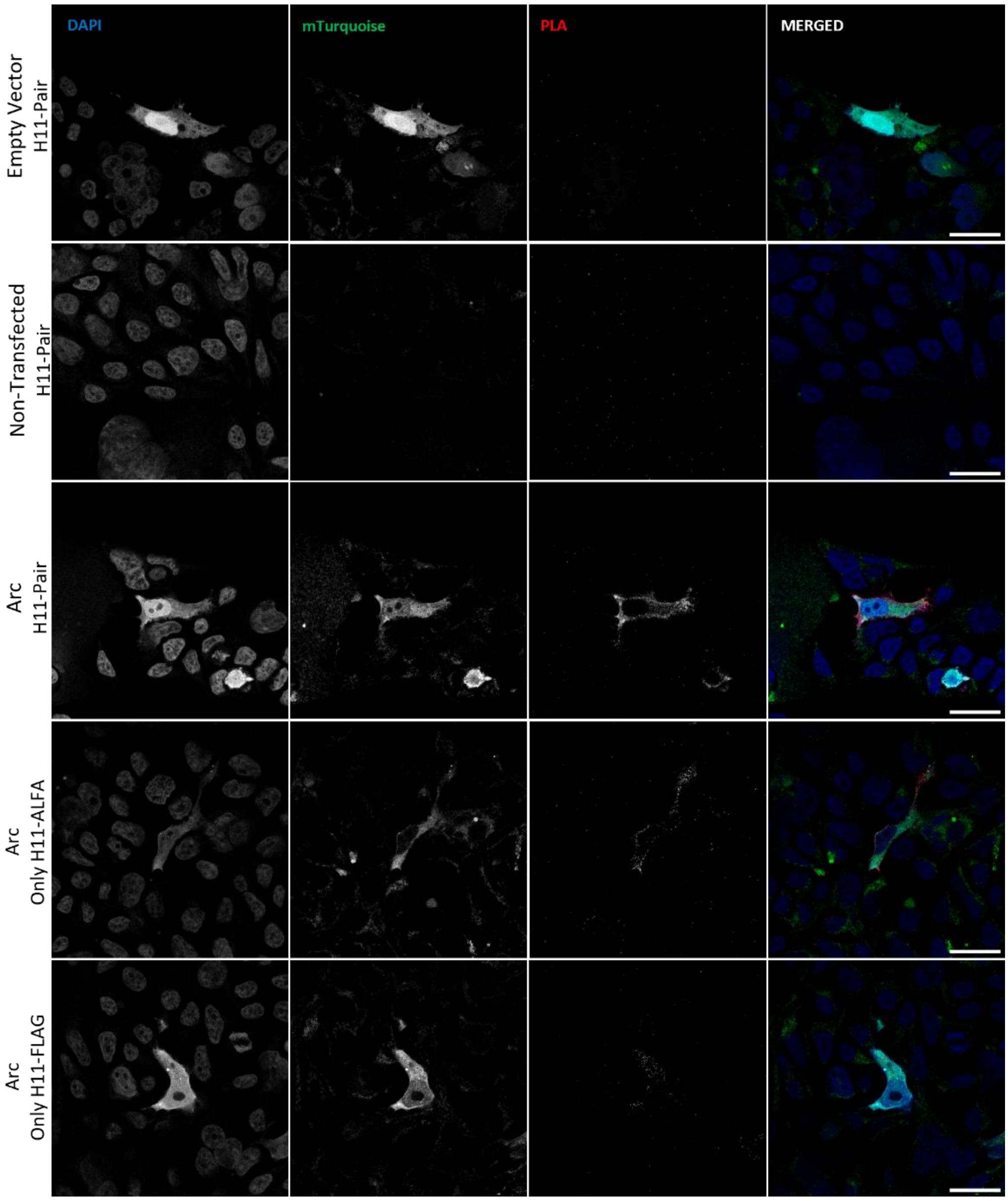


**Supplementary Figure 2. Arc-Arc nanobody-based PLA in HEK293FT cells is conditioned to the presence of ectopically expressed Arc protein.** Additional specificity controls for the validation of Arc-Arc PLA, arranged in 5 groups: (1) Empty vector + H11 pair ($1^{st}$ row) and (2) non-transfected cells + H11 pair ($2^{nd}$ row) as negative controls for Arc expression; (3) Arc-mTurquoise (mTq2) + H11 pair as positive group for Arc-Arc PLA; (4) Arc- mTq2 + only H11-ALFA or (5) H11-FLAG separately as negative controls for the H11 pair. DAPI staining the nucleus in blue, mTq2 as a transfection reporter in green and PLA signals in red. Scale bar of 25 µm.

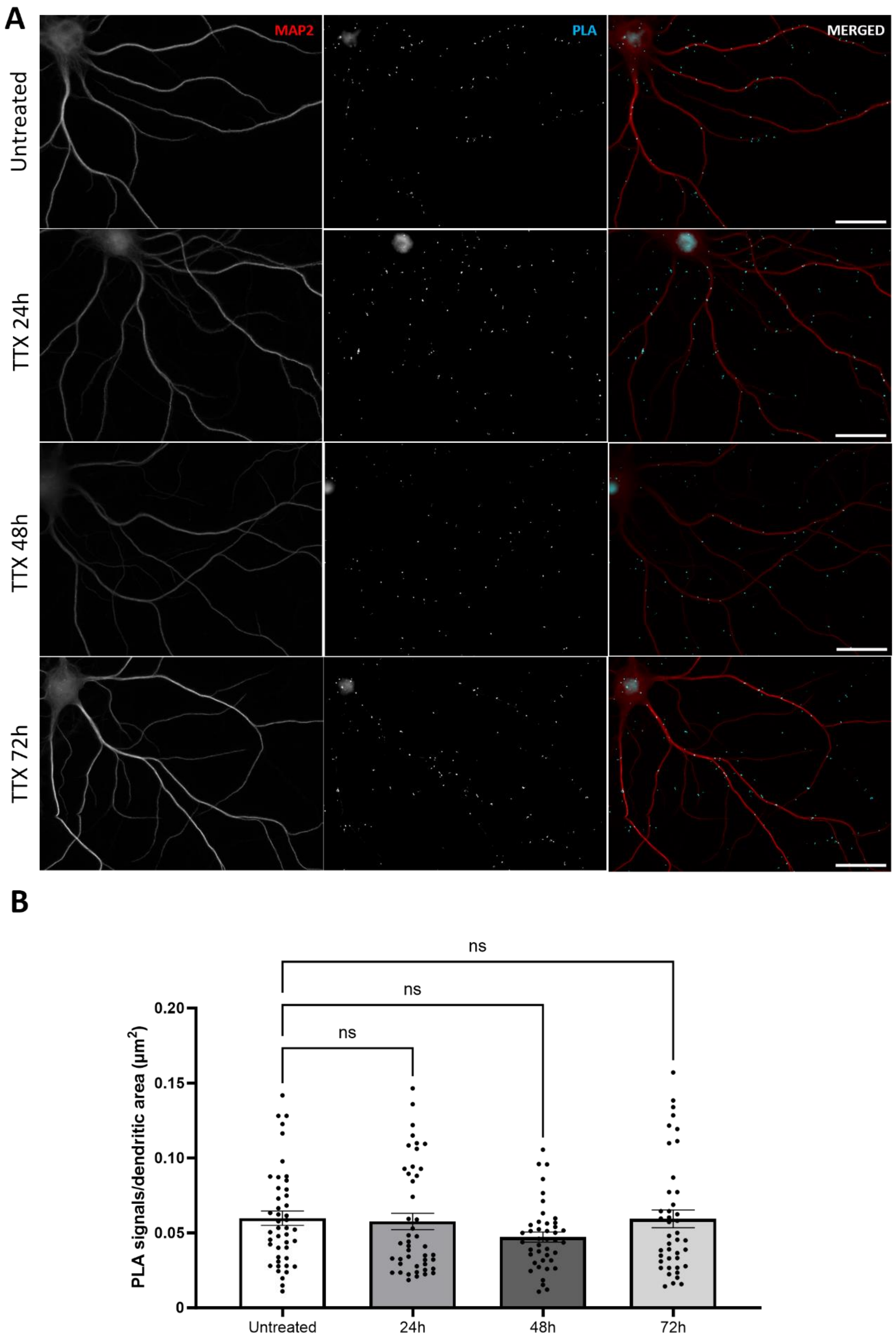


25

26 **Supplementary Figure 3. Longer TTX treatments (2 µM, 24-72h interval) do not affect Arc-**

27 **Arc PLA in hippocampal neuronal primary culture (DIV21) detected by H11 nanobody pair.**

**A** Representative image of treated and untreated groups. **B** Quantification of the comparison between treated groups and untreated groups, expressed in PLA signals/ dendritic area (µm2) ± SEM (45 images/group). One-way-ANOVA followed by Dunnett's multiple comparisons test showing p=0.9790, p=0.1853 and 0.9998, for 24, 48 and 72 hours TTX groups, respectively. Scale bar of 25 µm.

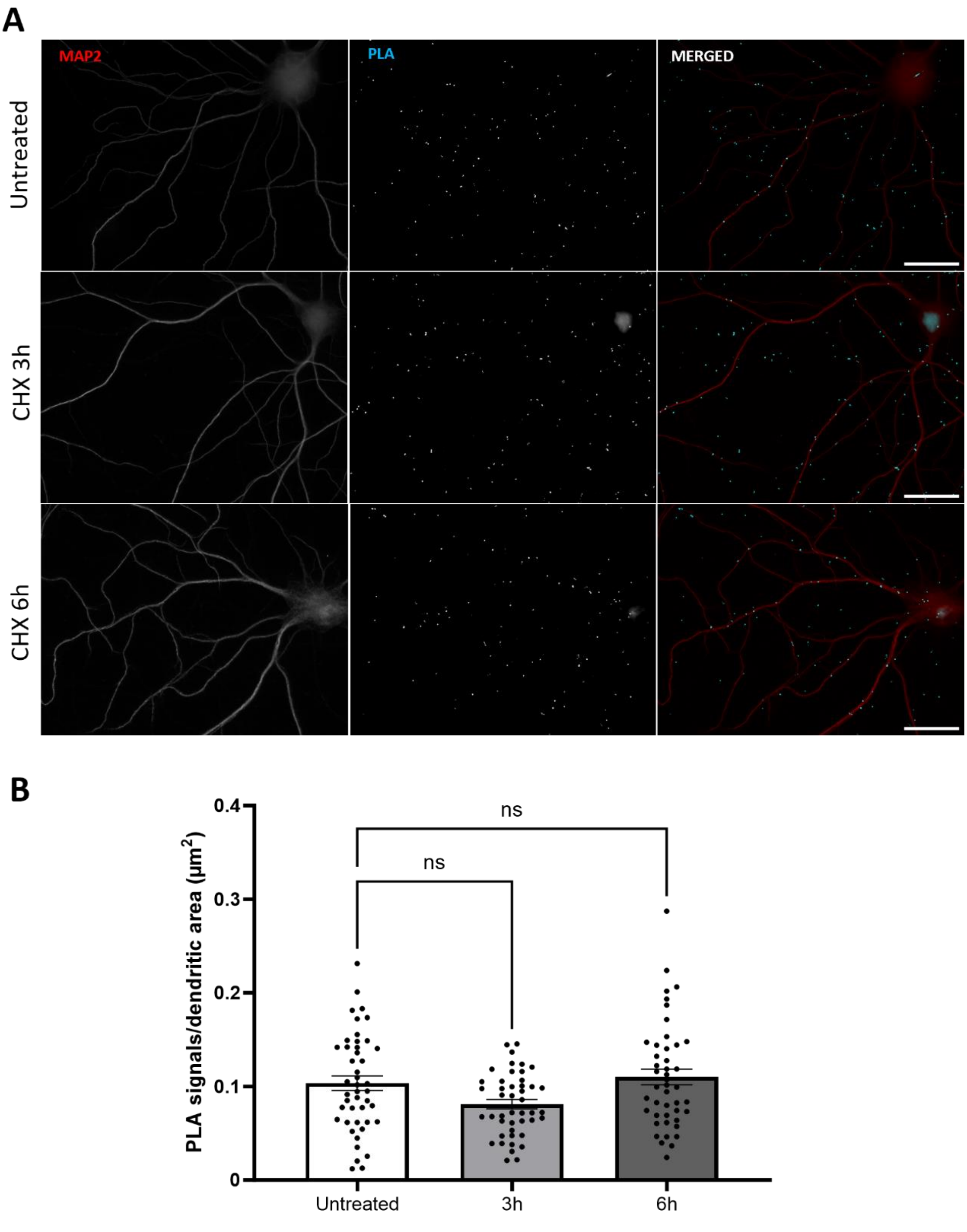


**Supplementary Figure 4. Longer CHX treatments (50 µg/mL, 3 and 6h interval) do not affect Arc-Arc PLA in hippocampal neuronal primary culture (DIV21) detected by H11 nanobody pair. A** Representative image of treated and untreated groups. **B** Quantification of the comparison between treated groups and untreated groups, expressed in PLA signals/ dendritic area (µm2) ± SEM (45 images/group). One-way-ANOVA followed by Dunnett's multiple comparisons test showing p=0.0546 and p=0.7342, for 3 and 6 hours CHX groups, respectively. Scale bar of 25 µm.

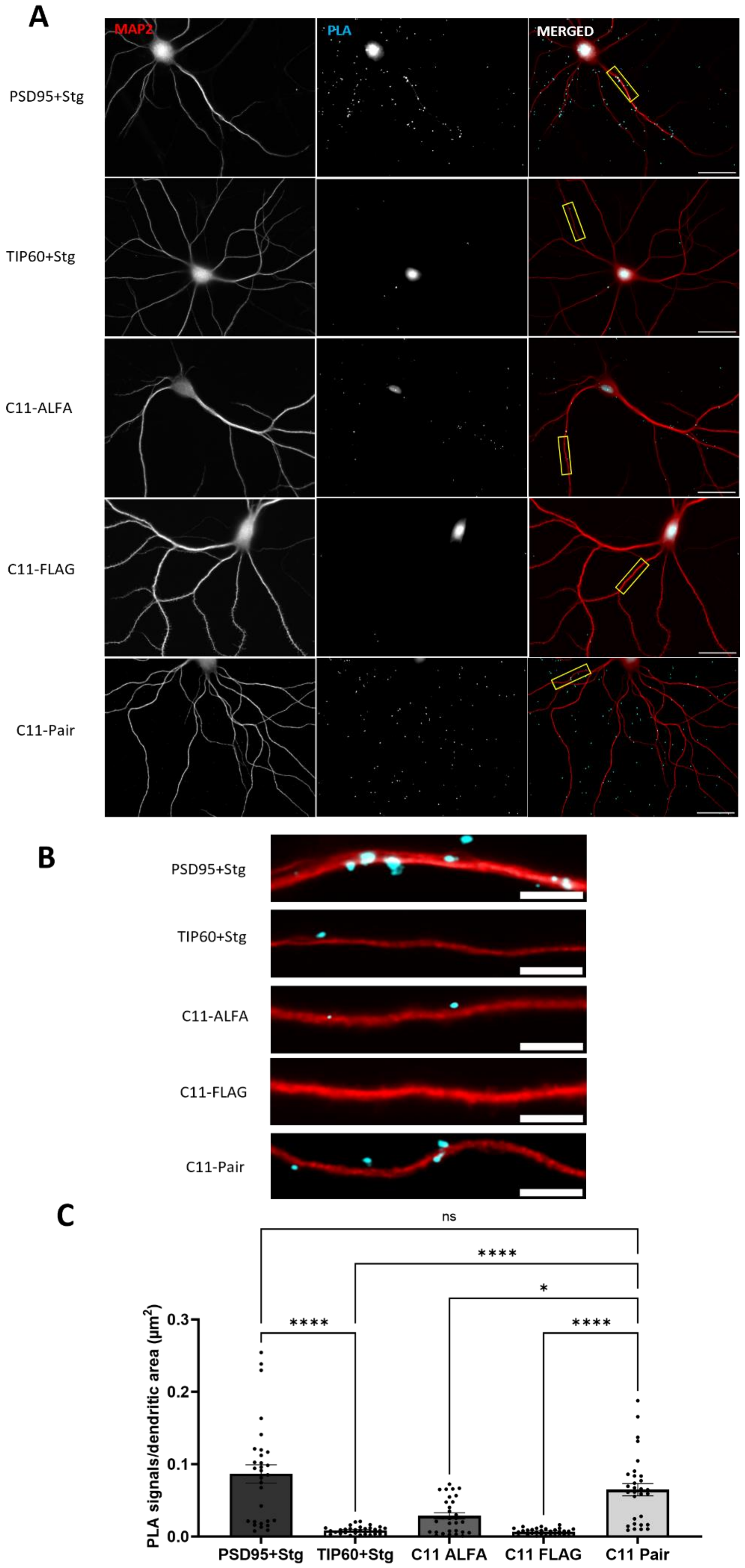
A
MAP2
PLA
MERGED
PSD95+Stg
TIP60+Stg
C11-ALFA
C11-FLAG
C11-Pair
B
PSD95+Stg
TIP60+Stg
C11-ALFA
C11-FLAG
C11-Pair
C
ns
****
****
*
****
PLA signals/dendritic area (µm²)
0.3
0.2
0.1
0.0
PSD95+Stg
TIP60+Stg
C11 ALFA
C11 FLAG
C11 Pair

43

**Supplementary Figure 5. Arc-Arc PLA validation for C11 nanobody pair in hippocampal neuronal primary culture (DIV21). A** Representative image from positive control (PSD95+Stg) and negative control (TIP60+Stg) antibody-based PLA; nanobody PLA using only ALFA-H11 or FLAG-H11 FLAG (rows 3 and 4, respectively), and nanobody PLA using the ALFA-H11/FLAG-H11 pair. Scale bar of 25 µm. **B** Digital zoom from corresponding groups in **A**. Scale bar: 5 µm. **C** Quantification of the method control expressed in PLA signals/ dendritic area ($\mu m^2$) ± SEM (30-45 images/group). One-way-ANOVA followed by Dunnett's multiple comparisons test showing *p≤0.05, **p≤0.01 and ****p≤0.0001.

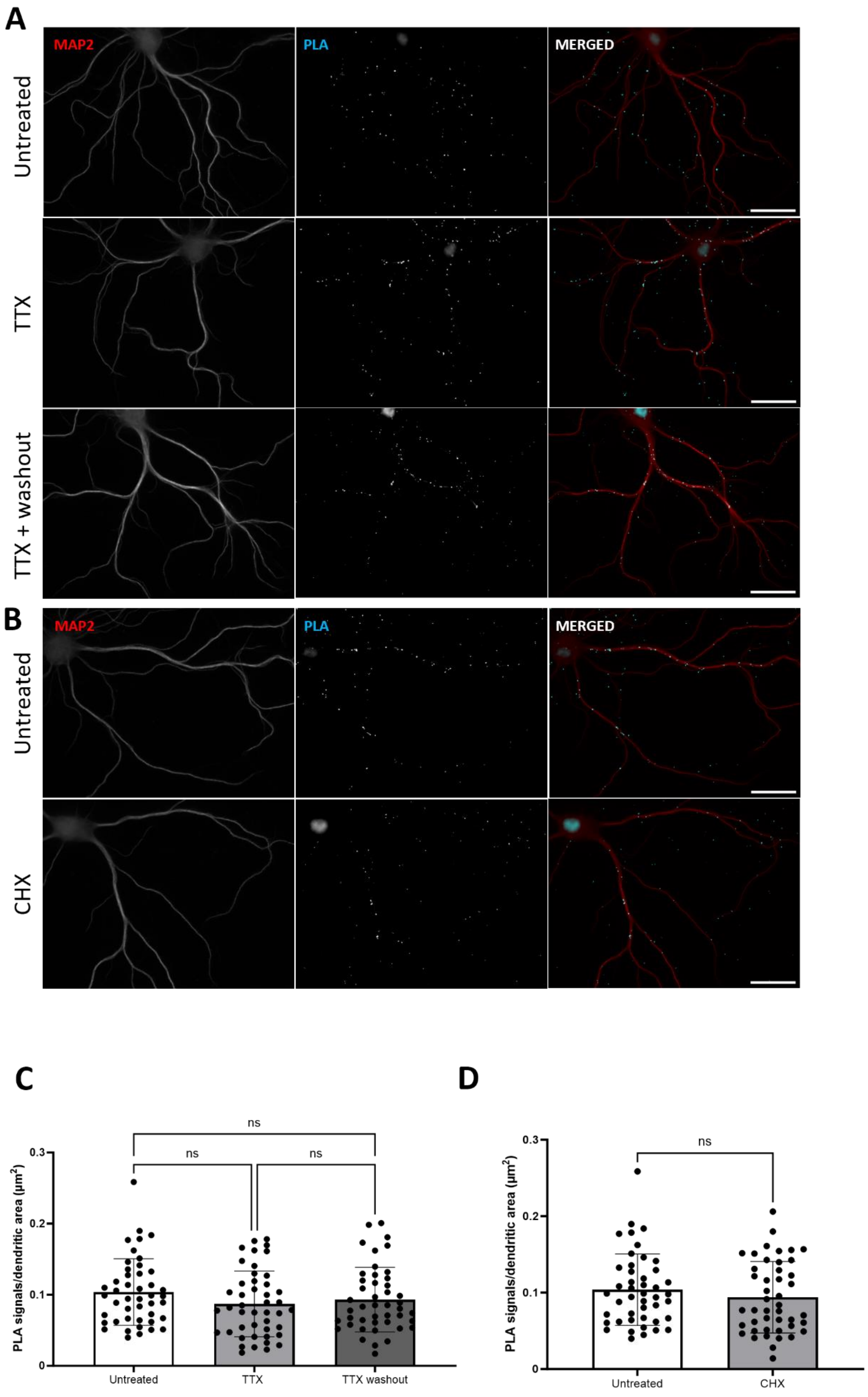
A
MAP2
PLA
MERGED
Untreated
TTX
TTX + washout
B
MAP2
PLA
MERGED
Untreated
CHX
C
ns
ns
ns
PLA signals/dendritic area (μm²)
0.3
0.2
0.1
0
Untreated
TTX
TTX washout
D
ns
PLA signals/dendritic area (μm²)
0.3
0.2
0.1
0
Untreated
CHX

**Supplementary Figure 6. Inhibition of neuronal firing by TTX and inhibition of protein synthesis do not affect basal Arc-Arc complexes detected by C11. A** Representative Arc-Arc C11 PLA images from hippocampal neuronal cultures treated with TTX for 16 h, TTX followed by 30 min washout, or no treatment. **B** Representative Arc-Arc C11 PLA images from hippocampal neuronal cultures treated with CHX for 70 min or no treatment. **C** Quantification of the comparison between treated (TTX and TTX + washout, $2^{nd}$ and $3^{rd}$ rows, respectively) groups and untreated groups, expressed in PLA signals/ dendritic area (µm2) ± SEM (45 images/group). One-way-ANOVA followed by Dunnett's multiple comparisons test showing p=0.1999 and p=0.5114, respectively. **D** Quantification of the comparison between untreated group and CHX treated group, expressed in PLA signals/ dendritic area (µm2) ± SEM (45 images/group). Unpaired Student's T-test showing p=0.3197. Scale bar of 25 µm.
